# Analyzing postprandial metabolomics data using multiway models: A simulation study

**DOI:** 10.1101/2022.12.19.521154

**Authors:** Lu Li, Shi Yan, Barbara M. Bakker, Huub Hoefsloot, Bo Chawes, David Horner, Morten A. Rasmussen, Age K. Smilde, Evrim Acar

**Affiliations:** *Department of Data Science and Knowledge Discovery, Simula Metropolitan Center for Digital Engineering, Oslo, Norway; Laboratory of Pediatrics, Section Systems Medicine and Metabolic Signalling, Center for Liver, Digestive and Metabolic Disease, University of Groningen, University Medical Center Groningen, Groningen, The Netherlands; Swammerdam Institute for Life Sciences, University of Amsterdam, Amsterdam, The Netherlands; Copenhagen Prospective Studies on Asthma in Childhood (COPSAC), Herlev and Gentofte Hospital, University of Copenhagen, Copenhagen, Denmark; Department of Food Science, University of Copenhagen, Copenhagen, Denmark

**Keywords:** postprandial metabolomics data, time-resolved metabolomics data, tensor factorizations (multiway data analysis), CANDECOMP/PARAFAC (CP), Principal Component Analysis (PCA), meal challenge test, whole-body metabolic model

## Abstract

**Background:** Analysis of time-resolved postprandial metabolomics data can improve the understanding of metabolic mechanisms, potentially revealing biomarkers for early diagnosis of metabolic diseases and advancing precision nutrition and medicine. Postprandial metabolomics measurements at several time points from multiple subjects can be arranged as a *subjects* by *metabolites* by *time points* array. Traditional analysis methods are limited in terms of revealing subject groups, related metabolites, and temporal patterns simultaneously from such three-way data.

**Results:** We introduce an unsupervised multiway analysis approach based on the CANDECOMP/PARAFAC (CP) model for improved analysis of postprandial metabolomics data guided by a simulation study. Because of the lack of ground truth in real data, we generate simulated data using a comprehensive human metabolic model. This allows us to assess the performance of CP models in terms of revealing subject groups and underlying metabolic processes. We study three analysis approaches: analysis of *fasting-state* data using Principal Component Analysis, *T0-corrected* data (i.e., data corrected by subtracting fasting-state data) using a CP model and *full-dynamic* (i.e., full postprandial) data using CP. Through extensive simulations, we demonstrate that CP models capture meaningful and stable patterns from simulated meal challenge data, revealing underlying mechanisms and differences between diseased vs. healthy groups.

**Conclusions:** Our experiments show that it is crucial to analyze both *fasting-state* and *T0-corrected* data for understanding metabolic differences among subject groups. Depending on the nature of the subject group structure, the best group separation may be achieved by CP models of *T0-corrected* or *full-dynamic* data. This study introduces an improved analysis approach for postprandial metabolomics data while also shedding light on the debate about correcting baseline values in longitudinal data analysis.

## Background

Postprandial metabolomics data (also referred to as meal challenge test data) includes the metabolic transition from the fasting to the fed state. Analysis of such data can reveal the underlying biological processes and improve the understanding of metabolic mechanisms [1, 2]. Examples include the study of a high-fat meal response for over one thousand subjects, which revealed subclass patterns in lipoprotein metabolism [3]. Analysis of postprandial metabolomics data also holds the promise to reveal new biomarkers especially for cardiometabolic diseases [4, 5]. In addition to detecting biomarkers and advancing prediagnosis, studying the postprandial state has proven useful in designing personalized nutrition [6]. Because of the high variability between individuals in terms of their post-meal glucose and triglyceride levels, personalized dietary interventions can be considered to lower individual post-meal glucose, thus may help decrease the risk of diseases such as prediabetes [6, 7].

There are various methods available for analyzing meal challenge test data. These are mainly supervised approaches, i.e., assuming that the group information (e.g., labels such as healthy and diseased) is known and incorporated into the analysis. In the aforementioned and other related work, univariate analyses are the most used tools for analyzing postprandial metabolism, e.g., the repeated measures analysis of variance (ANOVA) method is used to test group differences [3, 5]; a linear mixed model (LMM), which adds the random effect in the analysis, is used to explore the time and group interaction effect [8]. Univariate methods are simple and powerful in terms of studying group differences; however, such analyses are performed per metabolite, thus unable to reveal the relation between metabolites. Since metabolites are interlinked via pathways and hold dependent or independent relations with other metabolites, multivariate methods are promising in terms of capturing the underlying patterns in the data [4, 9]. Supervised multivariate methods used to analyze metabolomics data from challenge tests often combine ANOVA or LMM with principal component analysis (PCA). Such methods include ANOVA-Simultaneous Component Analysis (ASCA)[10], ANOVA-PCA [11], ANOVA-Partial Least Squares (PLS) [12], and their extensions [13, 14]. While these methods are suitable for time-resolved data, they rely on a priori known group information.

When the goal is exploratory analysis to reveal unknown stratification of subjects such as subgroups among healthy or diseased subjects, the workhorse data analysis approaches are unsupervised methods based on PCA and clustering [9]. However, these approaches cannot exploit the three-way structure of postprandial metabolomics data. They rely on either measurements at one time point or summaries such as clusters of time profiles obtained using data averaged across subjects [2, 15]. An effective way to preserve the multiway nature of the data and extract the underlying patterns is to use tensor factorizations [16–18], which are extensions of matrix factorizations such as PCA to multiway arrays (also referred to as higher-order data sets).

In this paper, we arrange time-resolved metabolomics data as a three-way array with modes: *subjects*, *metabolites*, and *time*, and use the CANDECOMP/PARAFAC (CP) [19, 20] tensor model to reveal the underlying patterns. The CP model summarizes the three-way array as a sum of a small number of factor triplets as in Figure 1. Among various tensor factorization methods, we use the CP model because of its uniqueness properties [16, 21], which facilitate interpretability of the extracted factors. In addition, the CP model is less sensitive to noise due to the concise structure (parsimony) of the model [22]. The CP model has previously been mentioned as a promising analysis tool for meal challenge test data [4, 9], but no analysis results have been presented for the three-way data arranged as in Figure 1. Recently, the effectiveness of the CP model in terms of analyzing other types of longitudinal data has been studied, e.g., using gut microbiome data from infants studying microbial changes and how those relate to the birth mode [23], urine metabolomics data from newborns exploring the use of the CP model for compositional data [24] and simulated dynamic metabolomics data relying on small-scale metabolic pathway models [25]. Unlike previous studies, we provide a comprehensive study of the CP model for analyzing time-resolved metabolomics data, in particular, focusing on the human metabolism in response to a meal challenge test and studying metabolic changes from a baseline state to dynamic states.

**Fig. 1:**
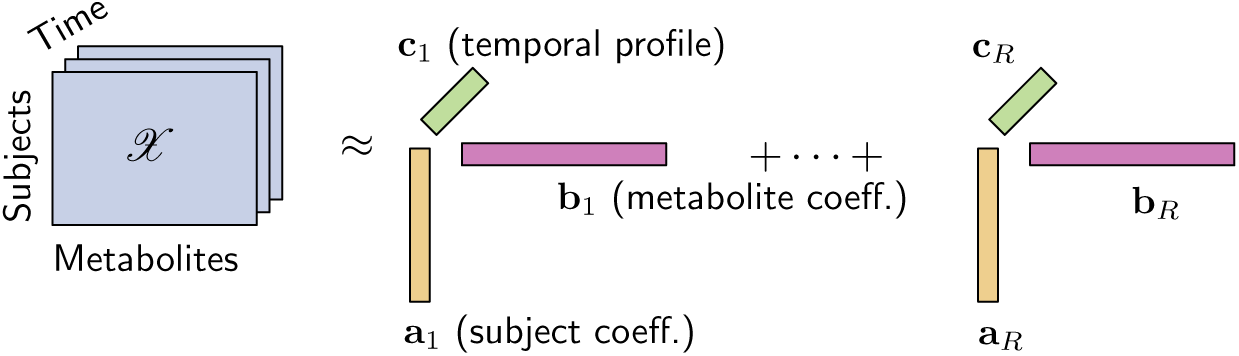
An *R*-component CP model of a three-way array with modes: *subjects*, *metabolites*, and *time*. Vectors a*_r_*, b*_r_* and c*_r_* correspond to the patterns in the *subjects*, *metabolites*, and *time* modes.

In the literature, analysis of *full-dynamic* data (i.e., full postprandial data without baseline correction) as well as *T0-corrected* data (i.e., the data corrected by subtracting the *fasting* state, similar to the method of analysis of changes [26]) have been considered [1, 2, 5, 27, 28]. Note that the *full-dynamic* data has information from the *fasting* as well as the *T0-corrected* (*pure-dynamic*) state. Understanding metabolic differences between these two states can help understand the metabolic mechanisms for postprandial metabolomics data. In this paper, we explore whether the CP model should be applied to *T0-corrected* data or *full-dynamic* data for postprandial metabolomics data analysis. Furthermore, we investigate the added value of analysis of the postprandial state compared to the fasting state.

In order to assess the performance of analysis approaches in terms of revealing the underlying patterns in the data, we generate simulated postprandial metabolomics data with known ground-truth information. To mimic the evolution of metabolite concentrations during a meal challenge, we used a human metabolic model [29] that describes metabolic pathways in eight organs in mechanistic detail by enzyme kinetic equations. Variation of disease-relevant parameters generated different subject groups (i.e., control vs. diseased). Through extensive computational experiments, we assess the performance of analysis of *fasting-state* data (using PCA), *T0-corrected* data (using a CP model) and *full-dynamic* data (using a CP model). We demonstrate that (i) CP models of postprandial metabolomics data reveal meaningful underlying patterns, (ii) for understanding metabolic differences among subject groups, it is crucial to analyze both *fasting-state* and *T0-corrected* data, (iii) depending on the nature of the subject group structure, the best performance in terms of revealing subject groups may be achieved by CP models of *T0-corrected* or *full-dynamic* data, and (iv) patterns extracted by CP models are reliable (i.e., consistently observed in a number of different settings such as when subsets of subjects are left-out, in the presence of a higher level of within-group variation, and with an unbalanced number of control and diseased subjects).

## Materials and methods

### Simulated postprandial metabolomics data

#### Human whole-body metabolic model

To generate simulated postprandial metabolomics data, we consider a human whole-body metabolic model proposed by Kurata [29]. The model is defined by a set of ordinary differential equations, with metabolites as variables and kinetic constants as parameters. Consisting of 202 metabolites, 217 reaction rates and 1,140 kinetic parameters, this multi-scale and multi-organ model is the largest, comprehensive, and highly predictive kinetic model of the whole-body metabolism [29]. It describes each reaction with a reversible Michaelis-Menten type rate equation and includes the action of insulin and glucagon, key hormones that govern the response to a meal (Figure 2). The meal in this simulated model from [29] includes 87 grams (g) carbohydrate and 33 g fat, and is given after an overnight fasting.

**Fig. 2:**
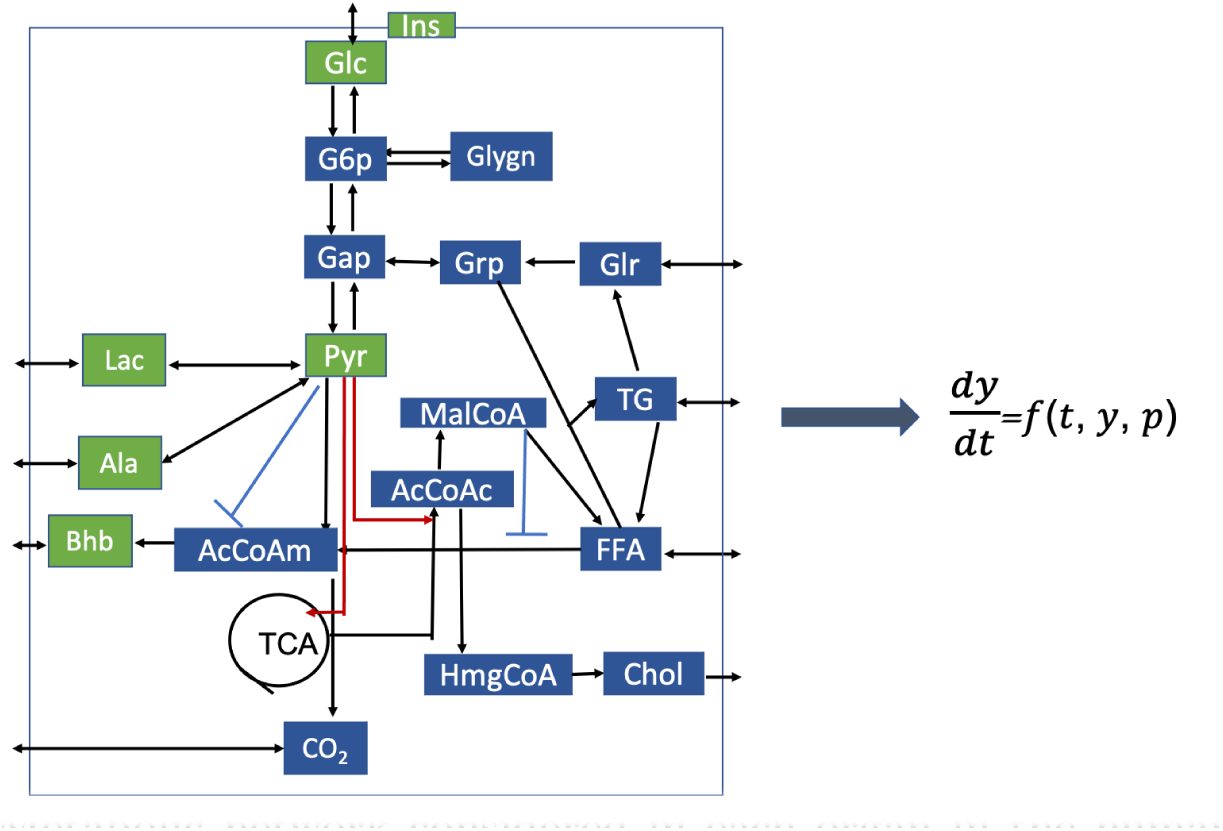
Metabolic network considered in each organ in the human whole-body model except for the pancreas; see [29] for more details.

#### Generation of the simulated data

To test whether our proposed unsupervised approach is able to reveal different subject groups, we generate simulated data using two types of variation: (i) between-group variation and (ii) within-group variation. The between-group variation is introduced by changing a specific parameter, i.e., Km Ins B M or Km inssyn Glc B, in the differential equations, as demonstrated in [29]. These two parameters are used to regulate the insulin-stimulated glucose uptake in skeletal muscle or the glucose-stimulated insulin secretion by the pancreas, respectively. Each type of between-group variation mimics one disease, which is as follows [29]:

- *Insulin resistance in skeletal muscle*: It is used for the study of type 2 diabetes mellitus and simulated by multiplying the default Km Ins B M with 1.5,
- *Beta-cell dysfunction*: Its extreme case is type 1 diabetes mellitus and it is simulated by multiplying the default Km inssyn Glc B with 1.1.

For the within-group variation, i.e., individual variation, we randomly perturb the kinetic constants in the liver (see Additional file 1 for a list of perturbed parameters). The perturbation level is denoted by *α*. For example, *α* = 0.2 indicates that related kinetic constants are set to be random parameters ranging from (100 *−* 20)% to (100 + 20)% of their default values.

Based on these definitions of between and within-group variations, we generate data for each subject as follows:

1. Get the 10-hour *fasting* concentrations for each individual (one individual has one set of random kinetic constants) by running the human whole-body metabolic model with the default initial values of model variables in [29], with an exception of the initial concentrations for particular metabolites presented in the next paragraph;
2. Start the meal challenge, i.e., run the human whole-body model using each individual’s 10-hour *fasting* state, and take the concentrations of metabolites at time points *t* = [0, 0.25, 0.5, 1, 1.5, 2, 2.5, 4] hours (with *T*_0_ = 0h, *T*_1_ = 0.25h, *· · ·*, *T*_7_ = 4h).

We choose the above time points to match the time samples in the real data, which we use to assess how realistic the simulations are (see Section *Simulated post-prandial metabolomics data are realistic*). In step one, we set initial values of Insulin (Ins) and blood metabolites Glucose (Glc), Pyruvate (Pyr), Lactate (Lac), Alanine (Ala), *β*-hydroxybutyrate (Bhb), Triglyceride and total Cholesterol to the median of fasting concentrations in the real data. Although the metabolic model involves 202 metabolites (including hormones) in different organs, we only consider hormone Ins and blood metabolites Glc, Pyr, Lac, Ala and Bhb since they are also measured in the real data. In addition, they are involved in the same metabolic network (see Figure 2), which makes the comparison of real and simulated data easier. When simulating data for a diseased subject, we use the parameter values given for *insulin resistance* and *beta-cell dysfunction*; otherwise, default values for parameters Km Ins B M and Km inssyn Glc B are used to simulate data for control subjects.

The simulated data from multiple subjects is arranged as a three-way array (# subjects *×* 6 metabolites *×* # time points) as in Figure 1.

#### Simulated postprandial metabolomics data are realistic

We compare the simulated data with the real data to demonstrate how realistic the simulations are. We use the real data corresponding to Nuclear Magnetic Resonance (NMR) spectroscopy measurements of plasma samples collected during a meal challenge test from the COPSAC_2000_ cohort [30]. The real data is not publicly available but may be shared by COPSAC through a collaboration agreement. The study was conducted in accordance with the Declaration of Helsinki and was approved by The Copenhagen Ethics Committee (KF 01-289/96 and H-16039498) and The Danish Data Protection Agency (2008-41-1754). Informed consent was obtained from the participants. The cohort considered in this work consists of 299 healthy 18-year-old subjects (144 males and 155 females). The blood samples were collected at eight time points following an overnight fasting during the fasting and postprandial states, i.e., at *t* = [0, 0.25, 0.5, 1, 1.5, 2, 2.5, 4]h. Over two hundred features were measured. We select the six features mentioned in Section *Generation of the simulated data* since these are the ones available in both real and simulated data.

Time profiles of metabolites in simulated and real data match well for most metabolites except for Pyr and Lac, where we noticed deviations (See Additional file 2: Figure S.1). The simulated Pyr differed most from the standard reference range [0.04, 0.1]mmol*/*L. To get a better correspondence for the initial concentrations, we performed a sensitivity analysis on the model and tuned the parameters related to Pyr and Lac (see Additional file 2: Figure S.2). After parameter tuning, all initial (fasting) concentrations, except Lac, corresponded well between the real and simulated data (see Figure 3). For Lac, we note that also in [29] the measured concentration of Lac is higher than the simulated value (1 vs. 0.5 mM). In our subjects, the Lac concentration is even higher, suggesting a difference between the cohorts. We do not necessarily expect a perfect correspondence between the real and simulated data throughout the time course since the simulated meal (taken from [29]) differs from the real meal (see Additional file 2: Table S1). In addition, the large individual variability in the real data puts the discrepancy between the real and simulated data in perspective, as demonstrated in Figure 3. The simulated model is realistic for our study in the sense that we observe that responses of key metabolites upon the meal challenge are in the physiological range, which is what we are interested in. Changing the metabolic model to simulate a meal-intake similar to the real data is outside the scope of this study.

**Fig. 3:**
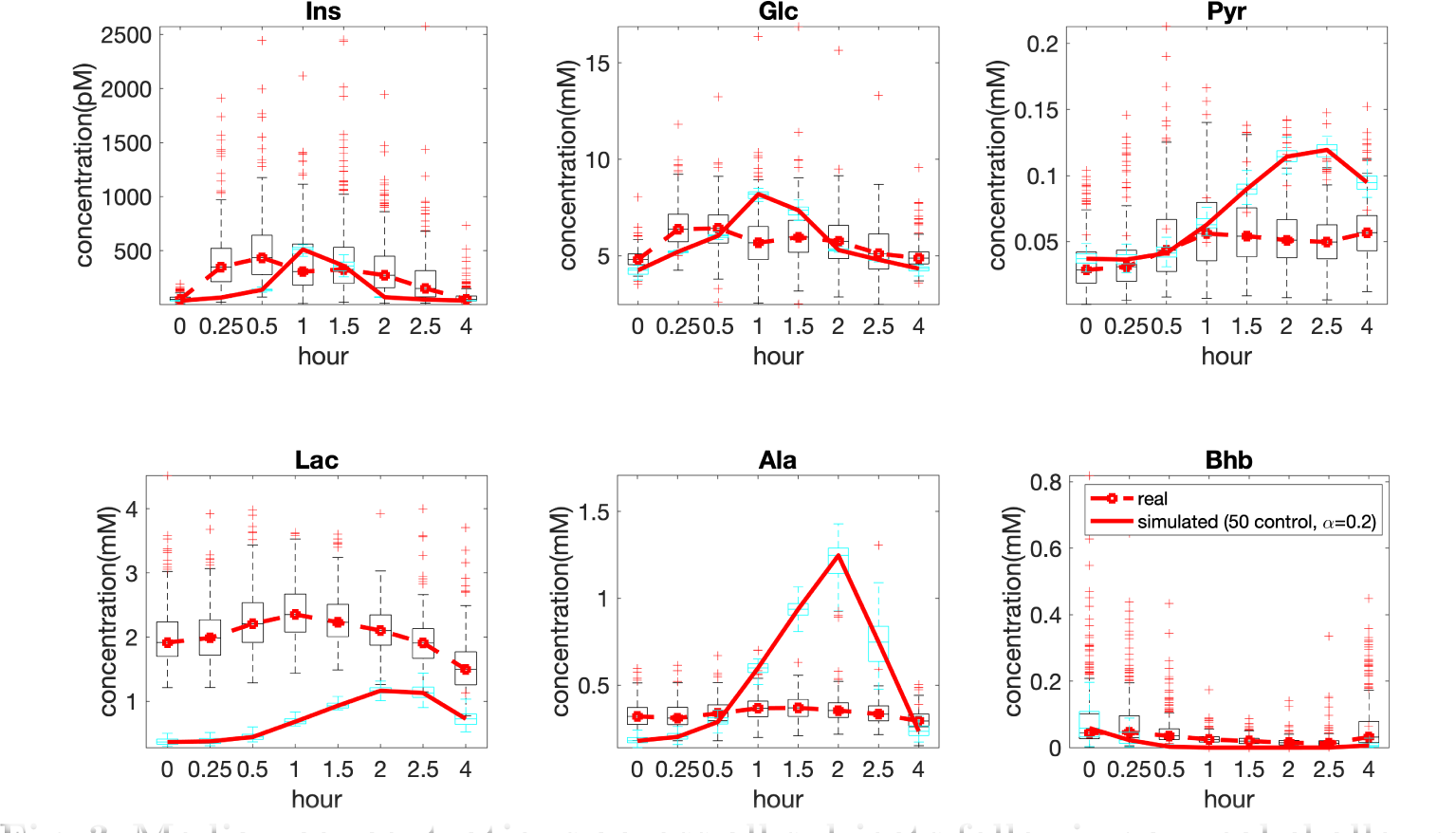
Median concentrations across all subjects following a meal challenge test. The standard meal in the real data includes 60 g palm olein, 75 g glucose and 20 g protein [31] and is given after overnight fasting, while the meal in simulations contains 87 g glucose and 33 g fat after 10-hour fasting.

### CANDECOMP/PARAFAC (CP) model

The CP model, which stands for Canonical Decomposition (CANDECOMP) [20] and Parallel Factor Analysis (PARAFAC) [19], stems from the polyadic form of a tensor [32]. Similar to the matrix Singular Value Decomposition (SVD), the main idea of the CP model is to represent a tensor as the sum of a minimum number of rank-one tensors (Figure 1). The CP model has been successfully used in many disciplines, including data mining [17, 33], neuroscience [34, 35], chemometrics [22], and recently also in longitudinal omics data analysis [23] in terms of revealing the underlying patterns from complex data sets.

For a third-order tensor *X ∈* R*^I×J×K^*, an *R*-component CP model represents the tensor as follows:

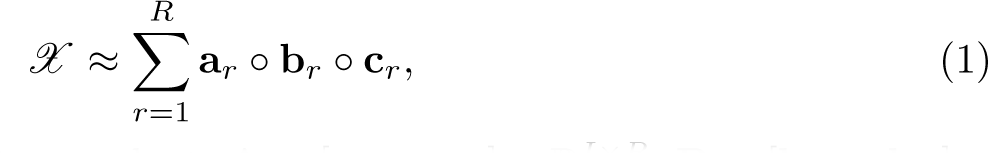

where *◦* denotes the vector outer product, **A** = [**a**_1_ *…* **a***_R_*] *∈* R*^I×R^*, **B** = [**b**_1_ *…* **b***_R_*] *∈* R*^J×R^*, and **C** = [**c**_1_ *…* **c***_R_*] *∈* R*^K×R^* are the factor matrices corresponding to each mode. Columns of the factor matrices, e.g., **a***_r_,* **b***_r_*, and **c***_r_* corresponding to the *rth component*, reveal the underlying patterns in the data. When assessing how well the CP model fits the data, we use the *explained variance* (also referred to as *model fit*) :

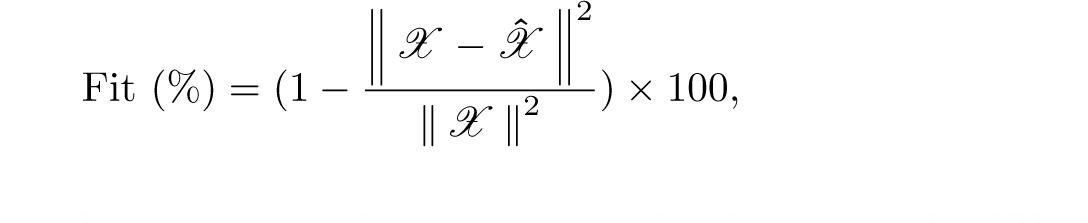

where 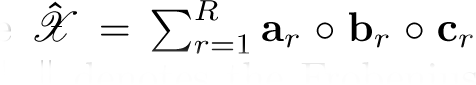 is the data approximation based on the CP model, and *∥ . ∥* denotes the Frobenius norm. A fit value close to 100% indicates that data *X* is explained well by the model; otherwise, there is an unexplained part left in the residuals.

The CP model has been widely used in applications as a result of its uniqueness property, i.e., factors are unique up to permutation and scaling ambiguities under mild conditions, without imposing additional constraints such as orthogonality [16, 21]. This uniqueness property facilitates interpretation. In the CP model, each rank-one component (**a***_r_,* **b***_r_,* **c***_r_*), i.e., *r*th column of factor matrices in all modes, can be interpreted, for instance, in terms of identifying groups of subjects (subjects with similar coefficients in **a***_r_*) positively or negatively related to specific sets of metabolites (metabolites with large coefficients in **b***_r_*) following certain temporal profiles (given by **c***_r_*). We normalize vectors (**a***_r_,* **b***_r_,* **c***_r_*) to unit norm when we interpret the factors. Due to the scaling ambiguity in the CP model, the norms can be absorbed arbitrarily by the vectors as long as the product of the norms of the vectors stays the same. In the presence of missing entries, which is common in applications, it is still possible to analyze such incomplete data using a CP model [36, 37].

The selection of the number of components (i.e., *R*) when fitting a CP model is a challenging task [16, 38], and an active research topic [39]. Among existing approaches, we make use of the model fit and the core consistency diagnostic [40]. In addition, we choose the number of components based on the interpretability of the model and replicability of the factors [39, 41] in subsets of the data (see Additional file 3 for details about the selection of number of components).

The performance of the CP model does not depend on the data size, i.e., the number of dimensions in each mode: *I*, *J*, *K*. If a sum of rank-one tensors, the CP model can reveal the underlying patterns even with data from a few subjects. For instance, fluorescence spectroscopy measurements of mixtures can be arranged as a third-order tensor with modes: mixtures, emission wavelengths, and excitation wavelengths. When such data is analyzed using a CP model, each rank-one component can capture one of the chemicals in the mixtures, and has done so successfully with only few, e.g., five, mixtures [22]. However, there are still challenges that will be encountered in the case of small data set size in any mode. First, the number of factors that can be uniquely extracted from the data will be limited [16, 21] as uniqueness conditions depend on the number of dimensions in each mode. Furthermore, using replicability to determine the number of components will not be possible with a small number of samples (that holds for a resampling strategy for any model). Finally, in the *time* mode, it is important to have enough time points to get temporal profiles.

### Analysis approach

We analyze each postprandial metabolomics data set using three methods: (i) PCA of the *fasting-state* data, (ii) CP model of the *T0-corrected* data, and (iii) CP model of the *full-dynamic* data. We compare the *fasting-state* analysis with *T0-corrected* analysis to understand metabolic differences between these two states. We compare these three approaches to obtain a better picture in terms of subject group differences and underlying metabolic mechanisms. When assessing subject group differences, we apply the unpaired (two-sample) *t*-test to each *subject* component and the null hypothesis is that the two groups come from independent random samples from normal distributions with equal means, without the assumption of equal variances. The main focus of the paper is to investigate metabolic differences associated with subject group differences during the *fasting* and *pure-dynamic* states, rather than striving for the best group separation.

### Experimental set-up and implementation details

In our experiments with simulated data, we consider the analysis of eight data sets generated using different types of between-group variation, different levels of individual variations, and balanced or unbalanced groups (see Table 1 for a complete list of data sets). In the balanced case, there are 50 controls and 50 diseased subjects; in the unbalanced case, there are 70 control and 30 diseased subjects.

**Table 1:** Eight simulated data sets generated using different settings. Here, α denotes the level of individual variation, where a smaller number indicates a lower level of individual variation. See the definition of *α* in Section *Generation of the simulated data*.

|  | Individual Variation |  |
| --- | --- | --- |
| *insulin resistance* | $\alpha = 0.2$ | balanced |
|  |  | unbalanced |
| | $\alpha = 0.4$ | balanced |
|  |  | unbalanced |
| *beta-cell dysfunction* | $\alpha = 0.2$ | balanced |
|  |  | unbalanced |
| | $\alpha = 0.4$ | balanced |
|  |  | unbalanced |

Before the CP analyses, the three-way data is preprocessed by centering across the *subjects* mode (to remove the common intercept from all subjects) and then scaling within the *metabolites* mode (to adjust for scale differences in different metabolites), i.e., dividing by the root mean squared value of each slice in the *metabolites* mode (see [42] for details about preprocessing multiway arrays). Before PCA, the *fasting-state* data is preprocessed similarly, i.e., centered and scaled.

For fitting CP models, we use cp-opt [43] and cp-wopt [37] from the Tensor Toolbox version 3.1 [44] with the Limited Memory BFGS optimization algorithm. cp-wopt uses weighted optimization [37] for fitting CP to incomplete data. For PCA, we use the svd function from MATLAB. Multiple random initializations are used to avoid local minima when fitting CP models. For the unpaired *t*-test, we use the ttest2 function from MATLAB.

When comparing CP models for different data sets, we use the factor match score (FMS) to quantify the similarity of factors in specific modes, e.g., the *metabolites* and *time* modes. FMS is defined as follows:

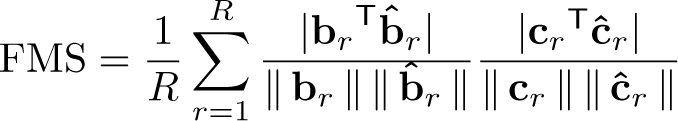

where **b***_r_*, **c***_r_* and **b^^^***_r_*, **^c***_r_* denote the *r*th column of factor matrices in the *metabolites* and *time* modes extracted by CP models from two different data sets (after finding the best permutation of the columns). FMS values are between 0 and 1, and an FMS value close to 1 indicates high similarity between the CP factors extracted from two different data sets.

All experiments are performed in MATLAB 2020a. Simulated data sets in Table 1 and example scripts are available in the GitHub repository for the paper: https://github.com/Lu-source/project-of-challenge-test-data.

## Results

### CP models extract similar factors from the simulated vs. real meal challenge data

Since subjects in the real data are considered healthy, we expect that the CP model extracts similar patterns in the *metabolites* and *time* modes from real data (299 subjects) vs. the simulated data with only the 50 control subjects (*α* = 0.2). Patterns extracted from the *subjects* mode are omitted since the *subjects* mode corresponds to different individuals in simulated vs. real data. Figure 4 shows that the 3-component CP model of each data set reveals similar patterns from the *metabolites* and *time* modes, to a certain extent.

**Fig. 4:**
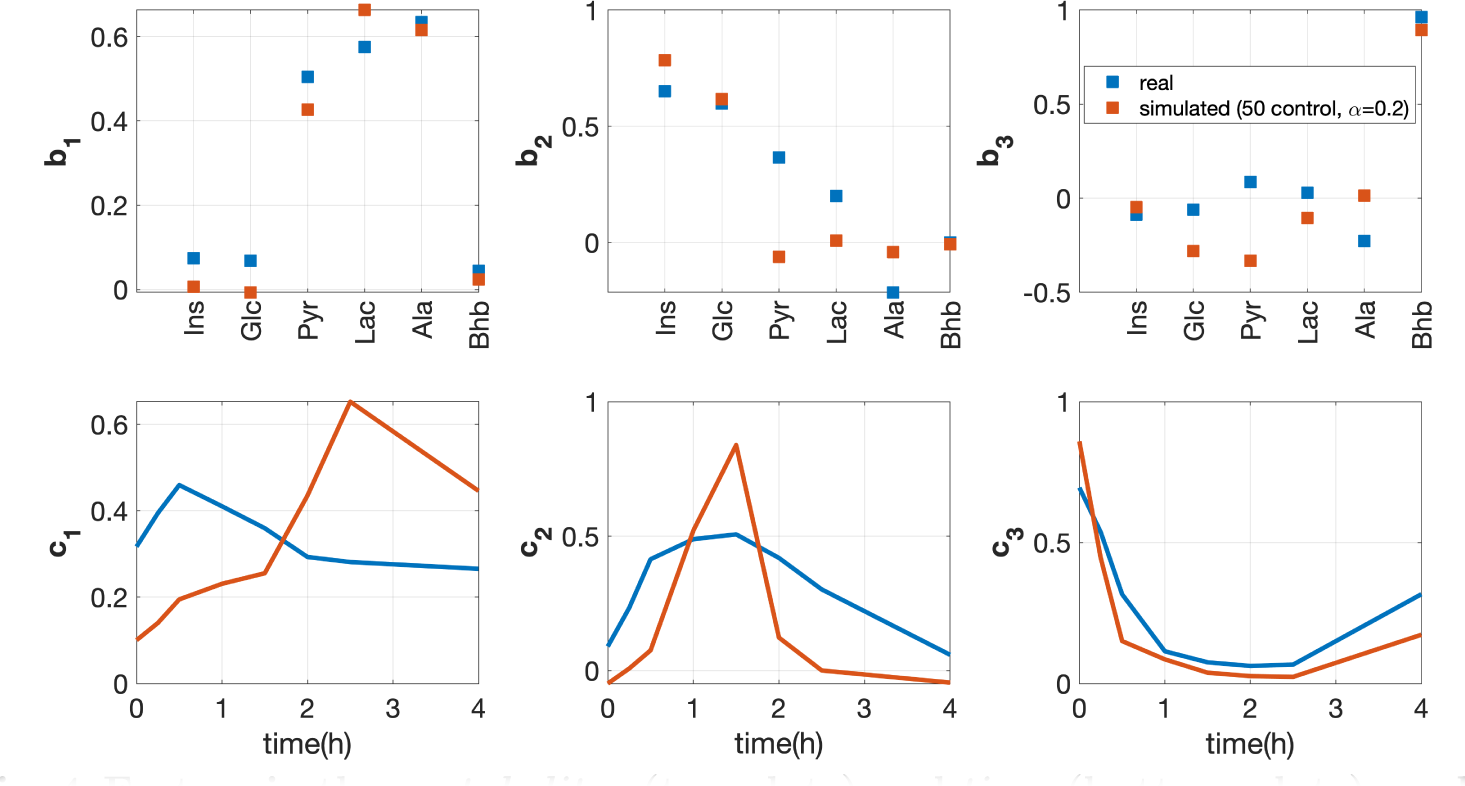
Factors in the *metabolites* (top plots) and *time* (bottom plots) modes extracted from the real vs. simulated data using a 3-component CP model. The vectors b**_1_**, b**_2_** and b**_3_** are the components in the *metabolites* mode, and vectors c**_1_**, c**_2_** and c**_3_** are the temporal patterns extracted by the CP model. Model fits for the real and simulated data are 50.3% and 71.6%, respectively.

The first component (**b**_1_ and **c**_1_) in Figure 4 mainly models metabolites Pyr, Lac and Ala (i.e., metabolites with large coefficients) which share similar dynamic patterns, i.e., being accumulated and then consumed. It makes sense that these three metabolites stay close since they are tied to each other with reactions Pyr*↔*Lac and Pyr*↔*Ala as shown in the pathway in Figure 2. However, we observe different peak heights and different time points for the highest peak in the *time* mode in real vs. simulated data. The different peak heights are possibly because meal compositions are different in real vs. simulated data. The different time points at the highest peak may be due to the fact that different individuals reach the highest peak at different time in the real data, and the first component of real data (with **c**_1_) captures the temporal profiles of a group of subjects with the highest peak appearing at around 0.5h in metabolites Lac, Ala and Pyr (see Additional file 4 for details).

The second component (see **b**_2_ and **c**_2_ in Figure 4) mainly captures Ins and Glc, which are accumulated and then consumed, after the meal challenge but at a different speed than Lac and Ala. However, we observe that the coefficient for Pyr is significantly different in real vs. simulated data. For the simulated data, the coefficient of Pyr is almost zero since the time point at the highest peak for all individuals in the simulated data is the same for each metabolite, and Pyr reaches to the highest peak at different time point compared with Ins and Glc. On the other hand, in real data, there are several subjects with dynamic patterns of Pyr reaching the highest peak at the same time point compared with Ins and Glc (see Additional file 4 for more details). This results in a positive coefficient for Pyr in **b**_2_ in real data.

The third component (**b**_3_ and **c**_3_ in Figure 4) mainly models Bhb, which behaves differently from other metabolites after a meal challenge, i.e., being consumed first and then getting accumulated. This behaviour has been observed in a consistent way in both real and simulated data.

### *Insulin-resistant* vs. control group with low within-group variation: *fasting* vs. *T0-corrected* analysis reveals metabolic differences and *T0-corrected* analysis performs better than *full-dynamic* analysis in terms of revealing group differences

#### Fasting-state analysis

PCA of the *fasting-state* data captures the subject group difference. The scatter plot of PC3 (the unpaired *t*-test using PC3 gives a *p*-value of 2 *×* 10*^−^*^9^) and PC5 (*p*-value of 2*×*10*^−^*^7^) in Figure 5 shows the best group separation. Note that PC3 and PC5 together explain only 21% of the data, indicating that the individual variation may dominate the total variation in the data. Figure 5 demonstrates that all metabolites except Ins and Glc are related to the subject group separation at the *fasting* state. Among them, Pyr contributes the most and is negatively related to the *insulin-resistant* group (i.e., the *insulin-resistant* group has mainly positive (negative) score values, and Pyr has a negative (positive) coefficient on PC3 (PC5)). This negative association indicates that the *insulin-resistant* group has lower concentration of Pyr at the *fasting* state. This results from a lower insulin-stimulated glucose uptake into the skeletal muscle in the *insulin-resistant* group, leading to less conversion of glucose into pyruvate.

**Fig. 5:**
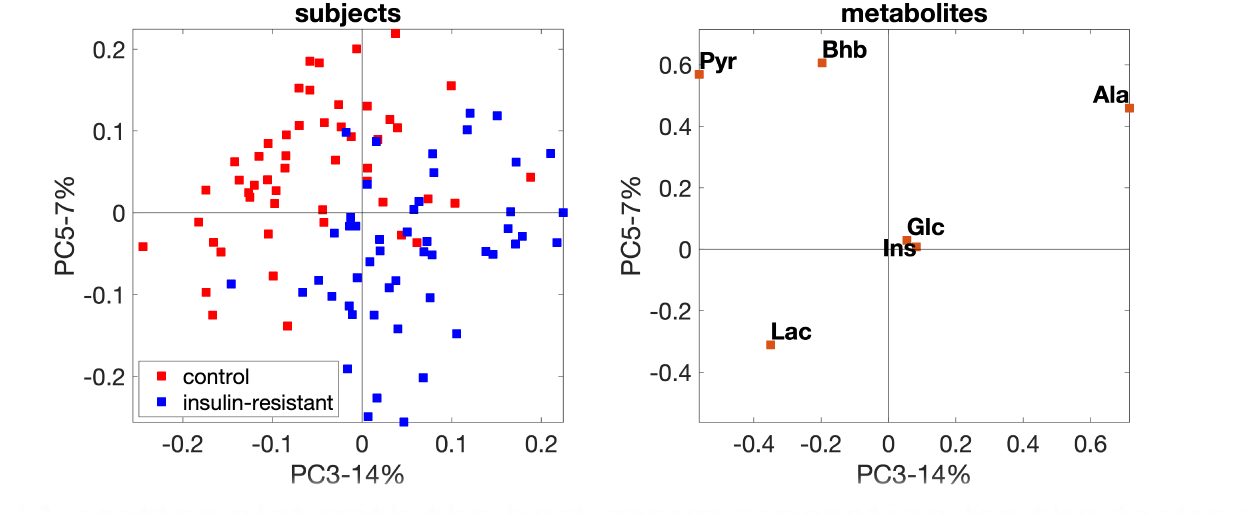
PCA scatter plot with the best group separation for the *fasting-state* data with *insulin-resistant* and control groups. The individual variation is introduced by setting the random perturbation level as ***α* = 0.2**.

#### T0-corrected analysis

We choose the 4-component CP model for the *T0-corrected* data (see Additional file 3 for the selection of number of components). This model reveals the subject group difference, as shown in Figure 6 (*subjects* mode). Among all components, the fourth component gives the best subject group separation (see the comparison of **a**_4_ with **a**_1_, **a**_2_ and **a**_3_). In the *metabolites* mode (see **b**_4_), metabolite Pyr has the largest coefficient, followed by Glc, while others have almost zero coefficients. In the *time* mode, **c**_4_ captures the dynamic behavior of mainly Pyr, i.e., it increases first and then slightly decreases. The increase of Pyr may be due to the glycolysis after the meal intake and its decrease afterwards may result from its conversion to other metabolites, e.g., Bhb, Lac and Ala, as shown in the pathway plot (Figure 2). **a**_4_, **b**_4_ and **c**_4_ together indicate that the concentration of Pyr increases more in the *insulin-resistant* group than in the control group; this suggests that the *T0-corrected* Pyr is positively related to the *insulin-resistant* group. Compared with **a**_4_, **a**_1_ and **a**_2_ reveal a weaker subject group separation due to the dynamic behavior of Lac & Ala (see **b**_1_ and **c**_1_) and Ins & Glc (see **b**_2_ and **c**_2_), respectively. Without capturing any group separation (see **a**_3_), the third component reveals that metabolite Bhb (it has the largest coefficient on **b**_3_ while others have almost zero coefficients) responds differently compared to other metabolites after the meal intake (see **c**_3_ vs. **c**_1_, **c**_2_ and **c**_4_), i.e., it decreases first and then slowly increases, as also shown in an earlier study [45].

**Fig. 6:**
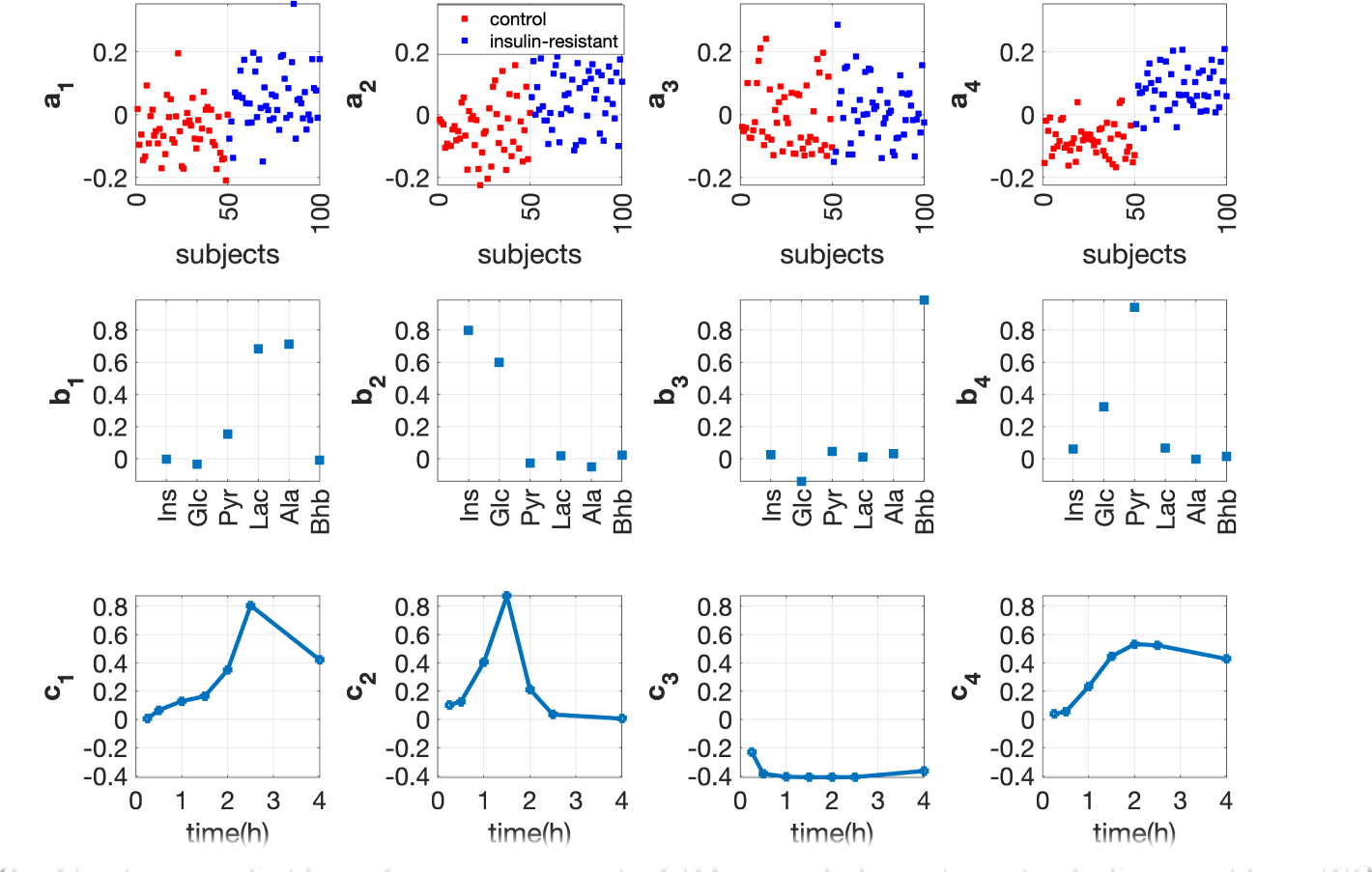
Factors of the 4-component CP model extracted from the *T0-corrected* data with *insulin-resistant* and control groups. The individual variation is set by using the random perturbation level ***α* = 0.2**. The vectors **a***_r_***, b***_r_* **and c***_r_*, *r* = 1*, · · ·,* 4**, are the components in the *subjects*, *metabolites*** and *time* modes extracted by the CP model.

#### Full-dynamic analysis

Figure 7 demonstrates the 4-component CP model of the *full-dynamic* data. This model captures both the *fasting-state* information (**c**_4_ reveals a constant temporal profile) and the *T0-corrected* information (i.e., the first and second components). The first component, i.e., **a**_1_, **b**_1_ and **c**_1_, and the second component, i.e., **a**_2_, **b**_2_ and **c**_2_, reveal the positive relation of Lac, Ala & Pyr and Ins & Glc with the *insulin-resistant* group and their dynamic response after the meal intake, respectively. The fourth component shows that Pyr negatively relates with the *insulin-resistant* group, i.e., the *insulin-resistant* group has negative coefficients in the *subjects* mode while Pyr has a positive coefficient in the *metabolites* mode.

**Fig. 7:**
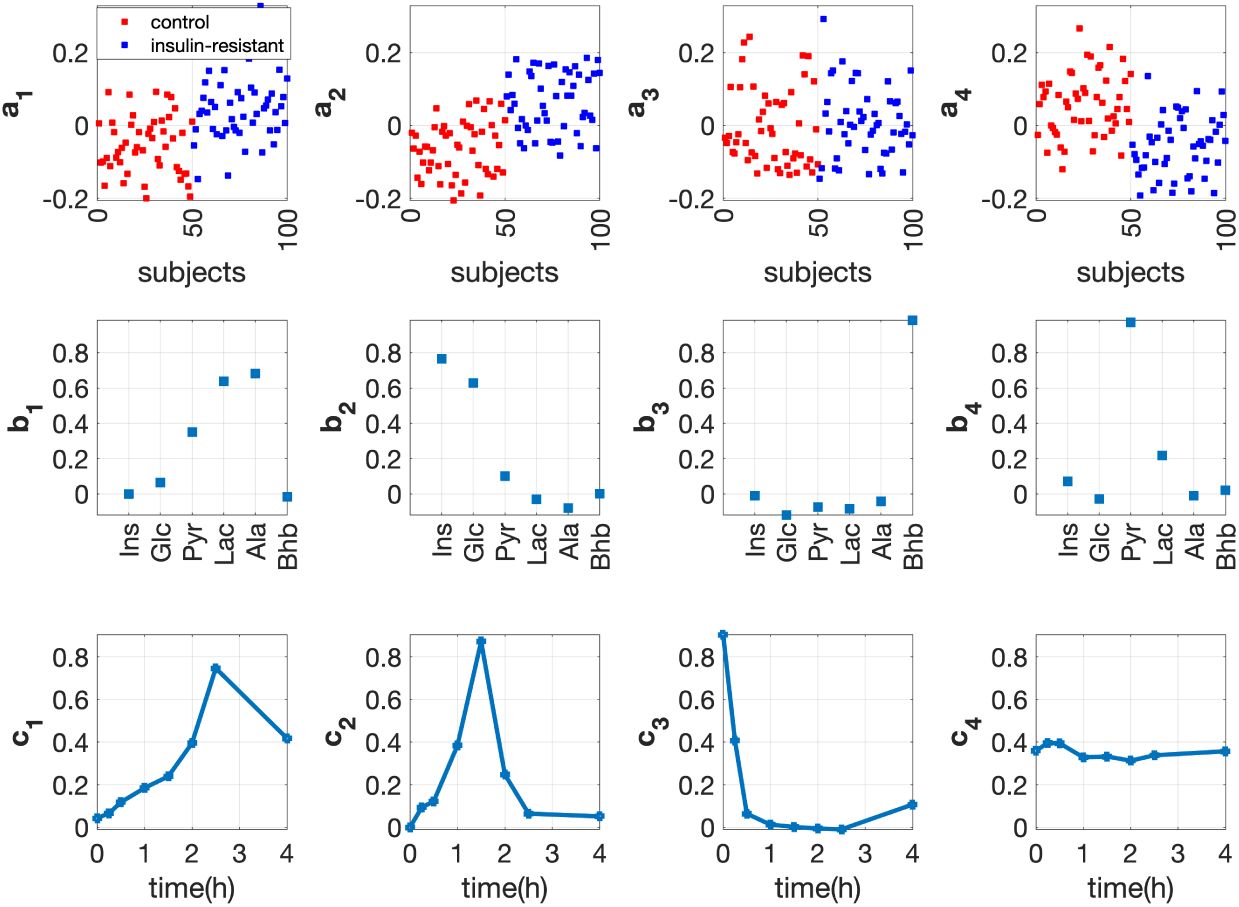
Factors of the 4-component CP model extracted from the *full-dynamic* data with *insulin-resistant* and control groups. The individual variation is set by using the random perturbation level ***α* = 0.2**. The vectors **a***_r_***, b***_r_* **and c***_r_*, *r* = 1*, · · ·,* 4**, are the components in the *subjects*, *metabolites*** and *time* modes extracted by the CP model.

#### Comparison of three analysis approaches

Through three analysis approaches, we observe four underlying mechanisms related to (i) Pyr, (ii) Ins & Glc, (iii) Lac, Ala & Pyr, and (iv) Bhb.

Pyr is identified by all methods as an essential metabolite related to the subject group separation. However, *fasting-state* vs. *T0-corrected* analysis reveals a different relation between Pyr and the *insulin-resistant* group. This, in turn, results in *T0-corrected* analysis performing better than *full-dynamic* analysis in terms of revealing group differences (see the comparison of the *subjects* mode in Figure 6 and 7). The best separation in *T0-corrected* analysis is given by the fourth component, where Pyr has the dominant coefficient in the *metabolites* mode. The left panel of Figure 8 shows that although the *insulin-resistant* group has a lower concentration of Pyr at the *fasting state*, Pyr accumulates much faster in the *insulin-resistant* group during the postprandial state (especially from 0.5h to 2h). In this sense, subtracting the *fasting-state* data from each time slice of the *full-dynamic* data leads to a more evident group difference between control vs. *insulin-resistant* subjects, as demonstrated in the right panel of Figure 8.

**Fig. 8:**
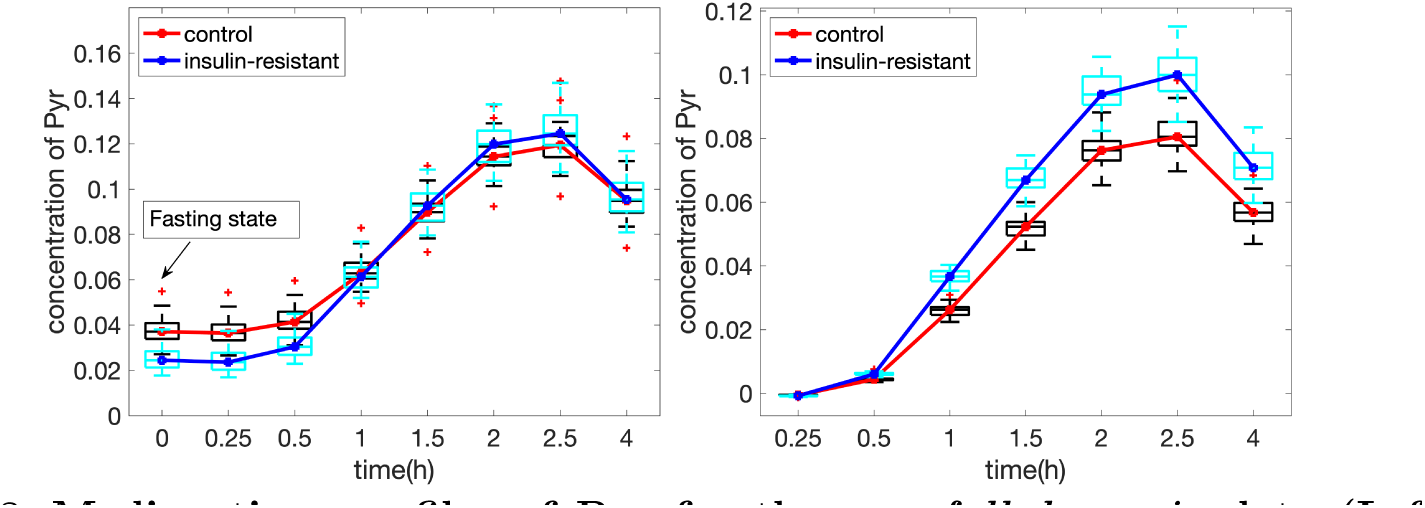
Median time profiles of Pyr for the raw *full-dynamic* data (Left panel) and *T0-corrected* data (Right panel) with *insulin-resistant* vs. control groups. The random perturbation level for generating individuals is *α* = 0.2.

The other mechanisms, (ii) Ins & Glc, (iii) Lac, Ala & Pyr, and (iv) Bhb, are captured through similar patterns in both *T0-corrected* and *full-dynamic* analysis.

### *Beta-cell dysfunction* vs. control group with low within-group variation: *fasting* vs. *T0-corrected* analysis reveals metabolic differences and *full-dynamic* analysis performs better than *T0-corrected* analysis in terms of separating subject groups

#### Fasting-state analysis

The *fasting-state* analysis captures the subject group difference. The scatter plot of PC3 (with *p*-value of 8 *×* 10*^−^*^7^ from the unpaired *t*-test) and PC5 (with *p*-value of 2 *×* 10*^−^*^8^) gives the best separation between the groups (Figure 9). Figure 9 also shows that metabolites Glc and Pyr contribute the most since they have large absolute coefficients along the discrimination direction (line *y* = *−x*). In addition, it is shown that metabolite Glc (Pyr) is positively (negatively) related to the *beta-cell dysfunction* group. This is consistent with Additional file 2: Figure S.3, which shows that the *beta-cell dysfunction* group has a higher concentration of Glc and a lower concentration of Pyr at the *fasting state* compared with the control group. The higher plasma Glc in the *beta-cell dysfunction* group results from inadequate glucose sensing to stimulate insulin secretion. Due to this inefficient secretion, glycolysis becomes less efficient, and less Pyr is produced, leading to a lower level of plasma Pyr in the *beta-cell dysfunction* group.

**Fig. 9:**
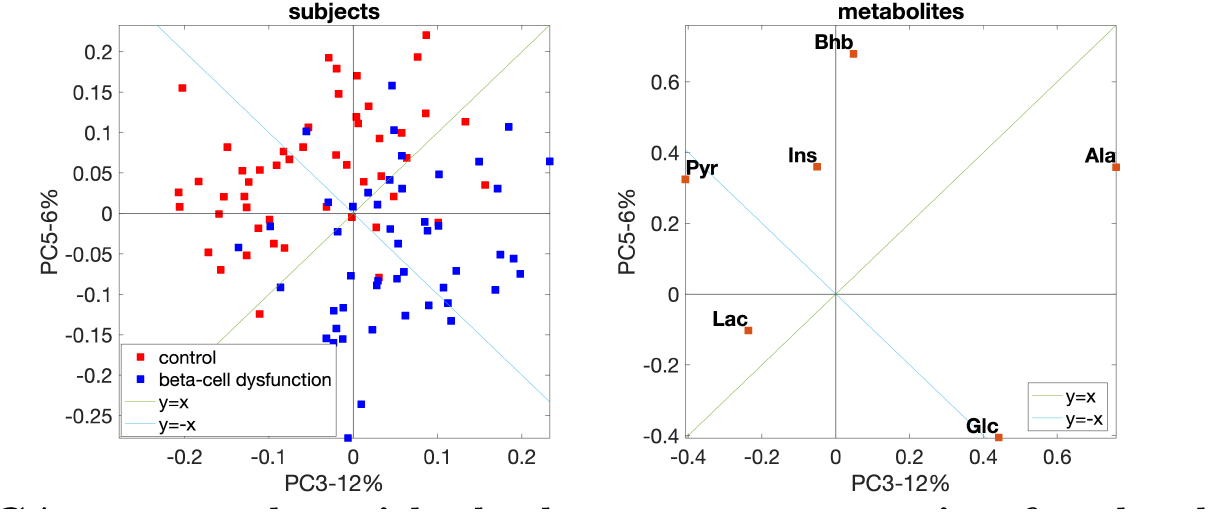
PCA scatter plot with the best group separation for the data set with *beta-cell dysfunction* and control groups. The individual variation is introduced by setting the random perturbation level to ***α* = 0.2**.

#### T0-corrected analysis

We choose the 4-component CP model for the *T0-corrected* data. This model captures the subject group separation, and the best separation is achieved by **a**_2_ (see Figure 10). In the *metabolites* mode (**b**_2_), Ins has the largest coefficient followed by Glc, while other metabolites almost have no contribution. This indicates that the dynamic changes of Ins and Glc contribute to the subject group separation, and Ins is the most significant one. Ins is shown to be negatively associated with the *beta-cell dysfunction* group. It makes sense that Glc stays close to Ins and shares a similar dynamic pattern as Ins (see Additional file 2: Figure S.4) since Ins and Glc regulate each other. In the *time* mode, **c**_2_ captures the dynamic pattern shared by Ins and Glc, i.e., an increase after the meal intake and then a decrease due to the glycolysis. **a**_1_ (with *p*-value of 8 *×* 10*^−^*^5^) and **a**_4_ (with *p*-value of 2 *×* 10*^−^*^4^) capture weak group separation in the *subjects* mode. In the *metabolites* mode, **b**_1_ captures the fact that Pyr, Lac and Ala respond to the meal challenge similarly, and this makes sense since they are close to each other, as shown in the pathway in Figure 2.

**Fig. 10:**
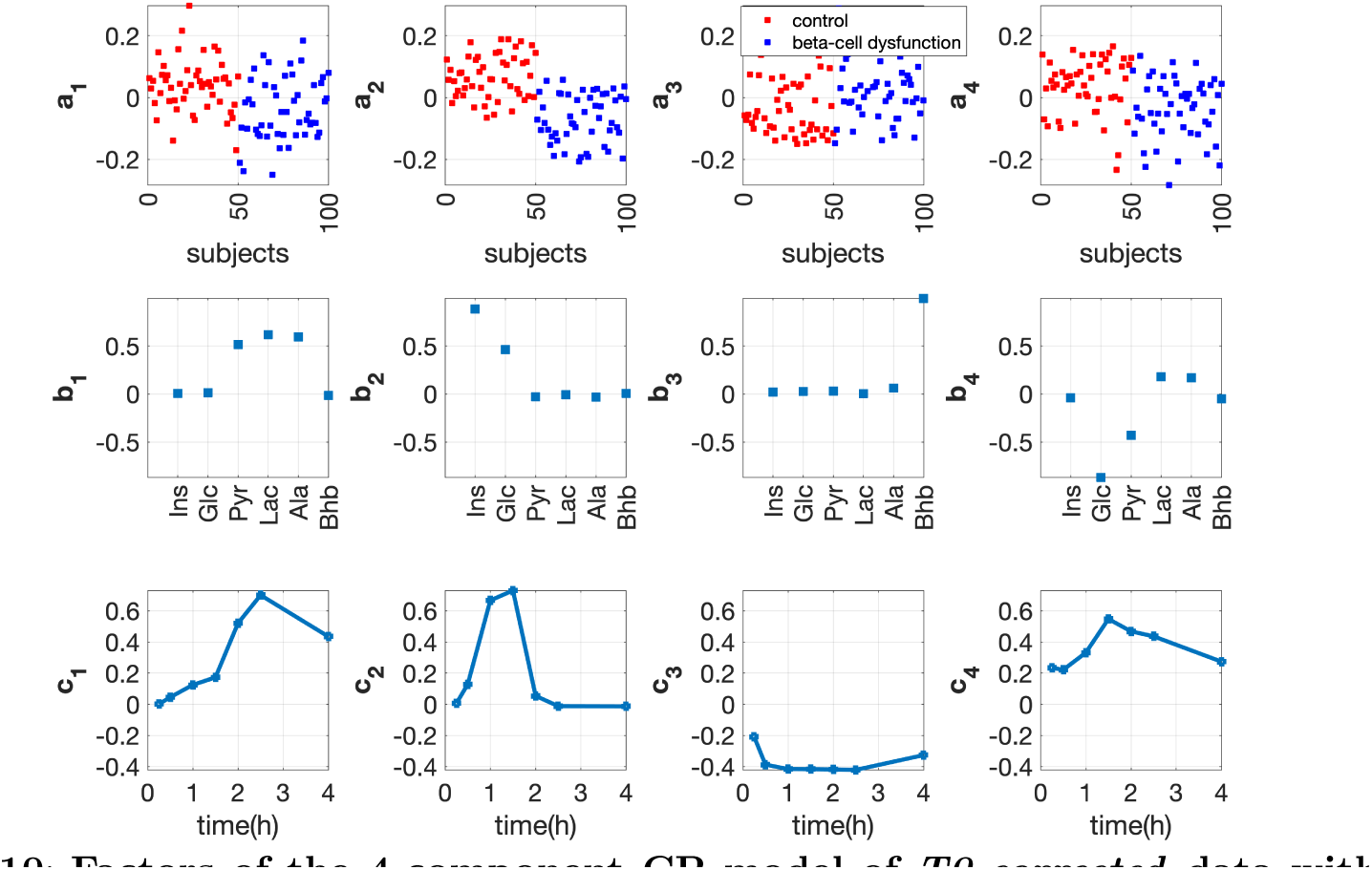
Factors of the 4-component CP model of *T0-corrected* data with *beta-cell dysfunction* and control groups. The individual variation is given by setting the random perturbation level to *α* = 0.2. The vectors a*_r_*, b*_r_* and c*_r_*, *r* = 1*, · · ·,* 4, are the components in the *subjects*, *metabolites* and *time* modes extracted by the CP model.

#### Full-dynamic analysis

We choose the 4-component CP model to analyze the *full-dynamic* data, with factor plots shown in Figure 11. **a**_4_ reveals the subject group differences best. In the *metabolites* mode (**b**_4_), metabolites Glc and Pyr have the largest coefficients in terms of absolute values. This component also reveals the positive (negative) relation of metabolite Glc (Pyr) with the *beta-cell dysfunction* group (i.e., the *beta-cell dysfunction* group has negative coefficients in the *subjects* mode while metabolite Glc (Pyr) has a positive (negative) coefficient in the *metabolites* mode). In the *time* mode, we observe a non-constant dynamic profile (**c**_4_) with a large coefficient at *t* = 0 comparable to the coefficients of other time points. This indicates that the fourth component captures a mixture of information from the *fasting* and *T0-corrected* states, and the *fasting-state* signal is relatively strong. The model also captures the pattern (the second component) in *T0-corrected* analysis (see **a**_2_, **b**_2_ and **c**_2_ in Figure 11), which shows that the beta-cell dysfunction group is negatively related with Ins.

**Fig. 11:**
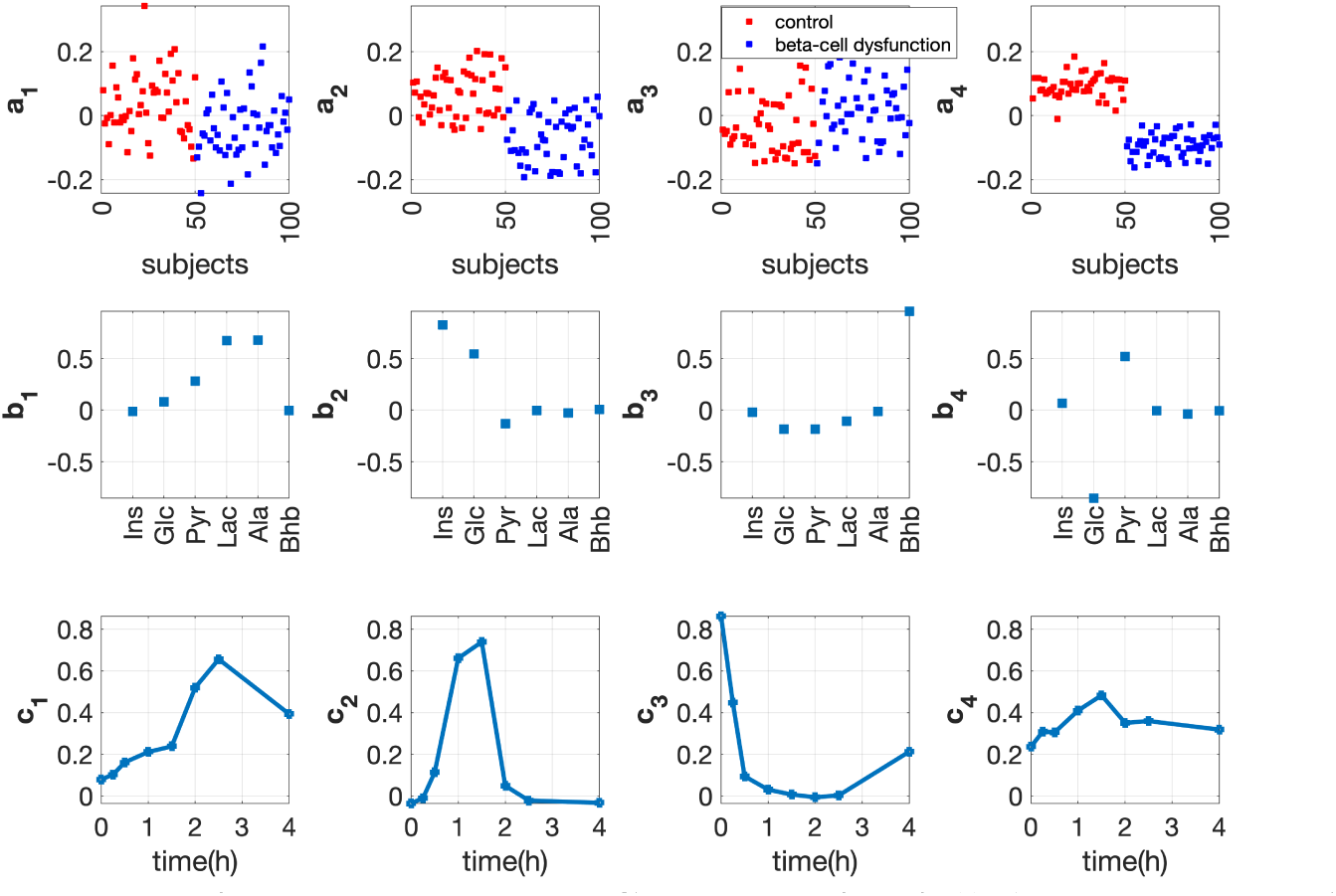
Factors of the 4-component CP model for *full-dynamic* data with *beta-cell dysfunction* and control groups. The individual variation is given by setting the random perturbation level to *α* = 0.2. The vectors a*_r_*, b*_r_* and c*_r_*, *r* = 1*, · · ·,* 4, are the components in the *subjects*, *metabolites* and *time* modes extracted by the CP model.

#### Comparison of three analysis approaches

In this case, we observe four underlying mechanisms related to (i) Glc & Pyr, (ii) Ins & Glc, (iii) Lac, Ala & Pyr, and (iv) Bhb.

While both *fasting-state* analysis and *T0-corrected* analysis can separate subject groups, the mechanisms responsible for group separation are different, with Glc & Pyr playing a role at the *fasting-state* analysis while Ins & Glc playing the main role in the *T0-corrected* analysis. We also observe that *full-dynamic* analysis outperforms the other two approaches in terms of capturing the subject groups (see the comparison of Figure 10 and 11). The best separation in the *full-dynamic* analysis is provided by the fourth component, which captures a mixture of the information from the *fasting* and *T0-corrected* states. The Glc & Pyr signal at the *fasting* state gets stronger through the dynamic information. This is consistent with the more evident group difference observed at the postprandial state from the time profiles of Glc (see Additional file 2: Figure S.3).

The other mechanisms (iii) Lac, Ala & Pyr, and (iv) Bhb are captured similarly using both *T0-corrected* and *full-dynamic* data analysis.

### The proposed analysis approach reveals metabolic aberrations

The crucial question in an unsupervised analysis is whether such an analysis can find groups among subjects pointing to metabolic differences or even metabolic deficiencies. The two studied metabolic deficiencies clearly show different patterns for the metabolites Ins, Glc and Pyr in the different analyses. These different patterns can be understood using the underlying metabolic network, thereby validating the power of the data analysis approach in terms of its ability to discriminate between different types of metabolic differences. More specifically, *insulin-resistant* subjects are expected to have higher *fasting* Glc levels, lower *fasting* Pyr levels, and higher *pure-dynamic* Ins levels. On the other hand, *beta-cell dysfunction* subjects are expected to have higher *fasting* Glc levels, lower *fasting* Pyr levels, and lower *pure-dynamic* Ins levels ([29] and references therein). The primary metabolic differences between these two metabolic deficiencies are related to Ins levels, with *insulin-resistant* group having a reduced ability to respond to Ins, resulting in higher Glc levels, which in turn stimulate further insulin secretion by the pancreas; conversely, those with *beta-cell dysfunction* have impaired insulin production, resulting in lower Ins levels. Figure 5 together with Figure 6, and Figure 9 together with Figure 10 indicate that our analyses can capture such metabolic differences between these two different types of metabolic deficiencies. Although we considered here only two metabolic deficiencies, we expect this property to hold also for other types of metabolic differences, especially when - in real data - there are more than six metabolites.

### CP models reveal stable patterns

Within-group variation can be enormous in real data making data analysis more challenging. For example, large within-group variations in healthy subjects might mask the changes of metabolites in response to diseases when exploring biomarker candidates [46]. Our numerical experiments demonstrate that CP models reveal stable patterns even when the level of individual variation is higher. Figure 12 shows the CP factors extracted using *T0-corrected* analysis from data sets generated with *insulin-resistant* and control groups using two different levels of individual variation, i.e., *α* = 0.2 and *α* = 0.4. We observe very similar factors in the *metabolites* and *time* modes even with different levels of individual variation. In addition, we consider data sets with an unbalanced number of control and diseased subjects. Table S.2 and Table S.3 in Additional file 2 show that CP factors are similar for data sets using balanced vs. unbalanced samples, and using different levels of individual variations, with the smallest FMS value of 0.90.

**Fig. 12:**
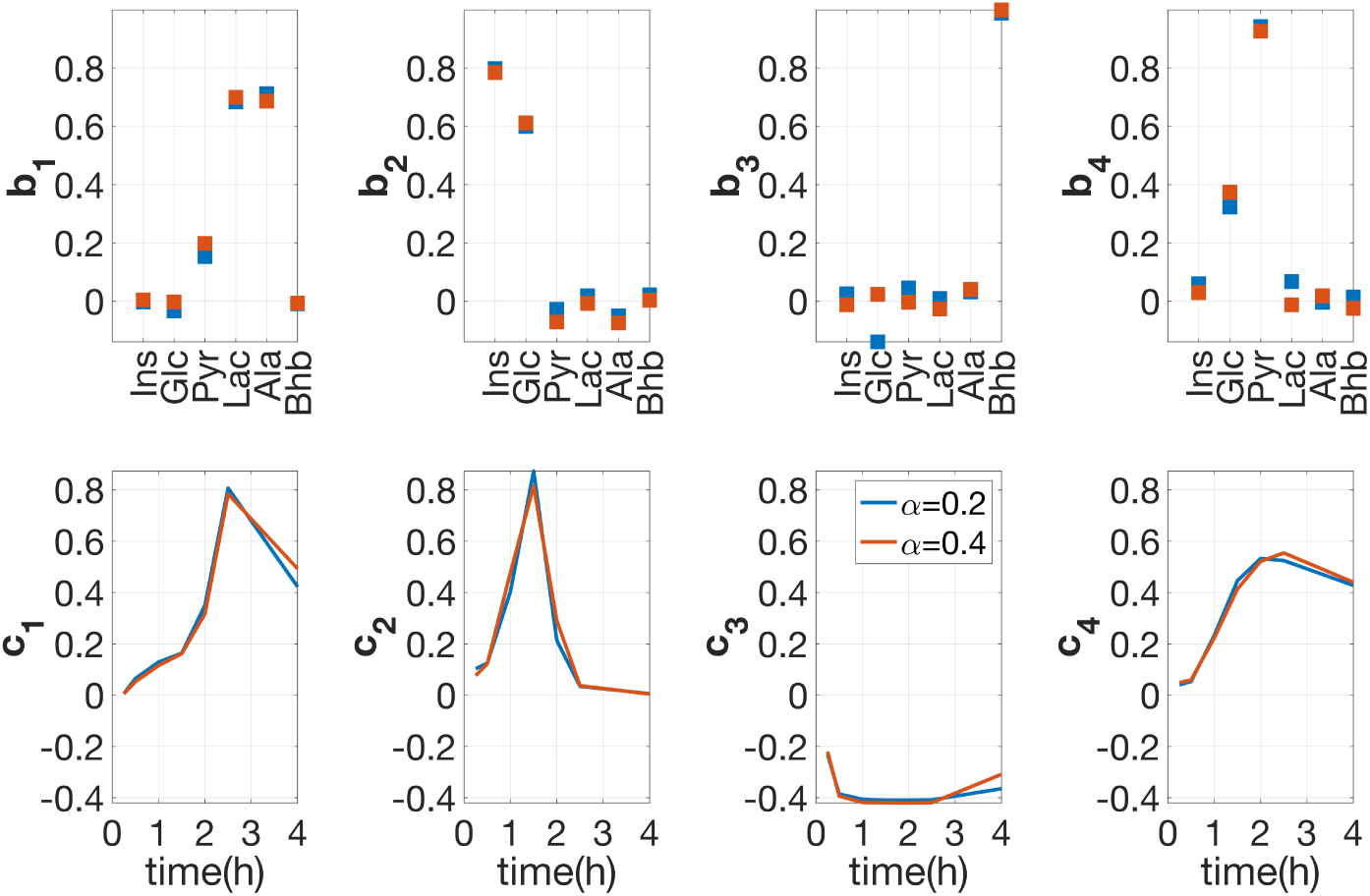
Comparison of the factors in the *metabolites* and *time* modes extracted from the *T0-corrected* data of two data sets with different levels of individual variation. Both data sets have 50 *insulin-resistant* subjects and 50 control subjects. Individual variations are introduced by setting the random perturbation level to ***α* = 0.2** in one data set vs. ***α* = 0.4** in the other **data set. The vectors a***_r_***, b***_r_* **and c***_r_*, *r* = 1*, · · ·,* 4**, correspond the components in the *subjects*, *metabolites* and *time* modes extracted by the CP models.**

The behaviours of CP models are also tested on data sets with random Gaussian noise mimicking the measurement error. The relative standard deviation (RSD) of the measurements encountered in this study can range from 4% to 15%, which aligns closely with the range mentioned in [47]. Our experimental findings indicate that the CP models are robust to noise when RSD of a replicate measurement is moderate. For example, when the RSD is 10% or 6% for the data set with *insulin-resistant* vs. control groups or the data set with *beta-cell dysfunction* vs. control groups, respectively, the deconvoluted CP patterns are very similar compared to their noise-free versions with the FMS values (noisy vs. noiseless patterns) greater than 0.92. With a further increase in RSD, the added noise may have an impact on the number of components in the CP models and the extracted patterns. More detailed results about comparisons of the patterns extracted by the CP models from noisy versus noiseless data can be found in Additional file 5.

## Discussion

In this paper, we have proposed an unsupervised approach based on the CP model to improve the analysis of the postprandial metabolomics data. The main focus is to demonstrate how the proposed method can effectively capture the differences in metabolic responses among different subject groups during a meal challenge test (in particular, a priori unknown stratifications rather than the setting where there are a priori labels as in, e.g., clinical interventions). Using the simulated data with known ground truth, we have demonstrated several benefits of the proposed approach: One benefit is the extraction of the underlying patterns in all modes simultaneously, which reveals subjects groups, related metabolites as well as corresponding temporal patterns from time-resolved postprandial data. In addition, we have shown that CP models can extract reliable patterns under various settings, e.g., even for data with considerable within-group variation. Another benefit is an overall picture of metabolic differences at the *fasting* vs. *T0-corrected* state, which can be obtained by using PCA for the *fasting-state* data and the CP model for the *T0-corrected* data analysis. Understanding such metabolic differences helps to explore the added value in the postprandial state due to metabolites’ dynamic responses. This, in turn, provides insightful guidance on why the *full-dynamic* analysis may capture a stronger or weaker group difference compared to the separate analyses of the *fasting-state* data and *T0-corrected* data in longitudinal analysis. The simulation studies indicate that such metabolic differences depend on the nature of the between-group variation, i.e., the data itself, which may be related to metabolic deficiency or aberration due to genetic variations in different groups of subjects. Therefore, to gain a more comprehensive understanding of the underlying metabolic mechanisms, particularly the interplay between the *fasting* and *T0-corrected* signals, we recommend using these three analyses together. Also, note that while our experiments rely on data sets consisting of a control and a diseased group, the proposed analysis approach holds the promise to reveal subject group differences due to different diseases, e.g., *insulin-resistant* vs. *beta-cell dysfunction* vs. control group, as well as subgroups of diseases, e.g., *insulin-resistant* vs. less *insulin-resistant* vs. control group. We expect the proposed approach to be useful also for other applications of time-resolved data analysis, for example, other types of challenge test data such as exercise challenge tests [48]. A common feature of such data is that they are time-resolved and hold both the baseline and *pure-dynamic* information. The essential idea is to analyze the baseline data together with the baseline-corrected data to understand their differences. If group structures are detected, understanding such differences will facilitate better utilization of the time-resolved information.

A side product of our study is an approach to generate meaningful simulated post-prandial metabolomics data. We use a comprehensive metabolic model to create such data. While the metabolic model was previously proposed, we tuned the model based on a comparison with our real data set including hundreds of subjects. Such simulated models may also be beneficial in terms of the optimal design of meal challenge tests. Nevertheless, there are also several limitations in this study. Using the metabolic model, we have only generated data where the concentration of each metabolite reaches the highest peak at the same time points for all subjects. However, in reality, time points at the highest peak vary from one subject to another. The main difficulty in generating such data is tuning the parameters while keeping the concentrations in reasonable ranges. Since the mathematical metabolic model involves 1140 parameters and 202 metabolites in eight different organs, parameter tuning is not trivial. Other limitations come from assumptions of the data analysis method. The CP model assumes that temporal patterns (i.e., **c***_r_*) are the same across subjects as we have in the simulated data; therefore, it may not be able to capture the individual differences such as shifted temporal profiles. Furthermore, another assumption in the CP model is that the factor matrix in the *metabolites* modes is the same across all time slices. However, those factor matrices may change over time. To account for such individual-specific temporal profiles or time-evolving factors in the *metabolites* mode, alternative multiway methods such as the PARAFAC2 model [49] may prove useful as in the analysis of neuroimaging signals [50–52] and in chemometrics [53]. We plan to study the performance of PARAFAC2 model in terms of analyzing dynamic metabolomics data as well as joint analysis of dynamic metabolomics data together with other omics measurements [54].

## Acknowledgments

This research has been funded by Research Council of Norway project 300489 and Novo Nordisk Foundation Grant NNF19OC0057934. We thank the children and families of the COPSAC_2000_ cohort for their contribution, and the clinical team at COPSAC for conducting the clinical testing.

## Supplementary information

### Additional file 1

A list of parameters perturbed for generating different individuals.

### Additional file 2

This file contains the following supplementary figures and tables. **Figure S.1.** Median time profiles of the real vs. simulated (50 control subjects and with the random perturbation level for individual variation set to *α* = 0.2) data generated from the default human whole-body metabolic model. **Figure S.2.** Sensitivity analysis for metabolites Lac (left panel) and Pyr (right panel). **Figure S.3.** Median time profiles of 50 control vs. 50 *beta-cell dysfunction* subjects for the *full-dynamic* data with individual variations introduced by setting the random perturbation level to *α* = 0.2. **Figure S.4.** Median time profiles of 50 control vs. 50 *beta-cell dysfunction* subjects for the *T0-corrected* data with individual variations introduced by setting the random perturbation level to *α* = 0.2. **Table S.1.** Meal compositions of the real and simulated data. **Table S.2.** *Full-dynamic data analysis*. Similarity between the CP factors (in the *metabolites* and *time* modes) extracted from different data sets and the CP factors of the data set with *α* = 0.2 and balanced samples. **Table S.3.** *T0-corrected data analysis*. Similarity between the CP factors (in the *metabolites* and *time* modes) extracted from different data sets and the CP factors of the data set with *α* = 0.2 and balanced samples.

### Additional file 3

Selection of the number of components in CP models.

### Additional file 4

Details about the comparison of the CP factors for the simulated vs. real data.

### Additional file 5

Experiment results for data sets with random noise.

## Declarations

### Ethics approval and consent to participate

All procedures performed in studies involving human participants were in accordance with the Declaration of Helsinki and was approved by The Copenhagen Ethics Committee (KF 01-289/96 and H-16039498) and The Danish Data Protection Agency (2008-41-1754). Informed consent was obtained from the participants.

### Consent for publication

Not applicable.

### Availability of code, data and materials

All code and simulated data are available in a GitHub repository at https://github.com/Lu-source/project-of-challenge-test-data. NMR measurements of plasma samples collected during the challenge test from the COPSAC2000 cohort are available through a collaboration agreement from COPSAC. For data access requests, please contact Morten A. Rasmussen.

## Authors’ contributions

A. K. S and E. A formulated the research problem. B. C and M. A. R prepared for the real data. B. M. B, H. H and L. L prepared for the simulated data. L. L performed the data analysis. S. Y and E. A validated the analysis. L. L, E. A, A. K. S, B. M. B and H. H interpreted the simulated data. L. L, E. A, A. K. S, B. M. B, H. H and D. H performed the comparison between the real and simulated data. L. L, A. K. S and E. A prepared the original draft. All authors were involved in the preparation of the final manuscript.

## References

[1] Harte, A.L., Varma, M.C., Tripathi, G., McGee, K.C., Al-Daghri, N.M., Al-Attas, O.S., Sabico, S., O’Hare, J.P., Ceriello, A., Saravanan, P., et al.: High fat intake leads to acute postprandial exposure to circulating endotoxin in type 2 diabetic subjects. Diabetes Care 35(2), 375–382 (2012)

[2] Wopereis, S., Stroeve, J.H.M., Stafleu, A., Bakker, G.C.M., Burggraaf, J., Erk, M.J., Pellis, L., Boessen, R., Kardinaal, A.A.F., Ommen, B.: Multi-parameter comparison of a standardized mixed meal tolerance test in healthy and type 2 diabetic subjects: the phenflex challenge. Genes & Nutrition 12(21), 1–14 (2017)

[3] Wojczynski, M.K., Glasser, S.P., Oberman, A., Kabagambe, E.K., Hopkins, P.N., Tsai, M.Y., Straka, R.J., Ordovas, J.M., Arnett, D.K.: High-fat meal effect on ldl, hdl, and vldl particle size and number in the genetics of lipid-lowering drugs and diet network (goldn): an interventional study. Lipids in Health and Disease 10(181), 1–11 (2011)

[4] Ĺepine, G., Tremblay-Franco, M., Bouder, S., Dimina, L., Fouillet, H., Mariotti, F., Polakof, S.: Investigating the postprandial metabolome after challenge tests to assess metabolic flexibility and dysregulations associated with cardiometabolic diseases. Nutrients 14(3), 472 (2022)

[5] Kumar, A.A., Satheesh, G., Vijayakumar, G., Chandran, M., Prabhu, P.R., Simon, L., Kutty, V.R., Kartha, C.C., Jaleel, A.: Postprandial metabolism is impaired in overweight normoglycemic young adults without family history of diabetes. Scientific Reports 10, 353 (2020)

[6] Zeevi, D., Korem, T., Zmora, N., Israeli, D., Rothschild, D., Weinberger, A., Ben-Yacov, O., Lador, D., Avnit-Sagi, T., Lotan-Pompan, M., et al.: Personalized nutrition by prediction of glycemic responses. Cell 163(5), 1079–1094 (2015)

[7] Berry, S.E., Valdes, A.M., Drew, D.A., Asnicar, F., Mazidi, M., Wolf, J., Capdevila, J., Hadjigeorgiou, G., Davies, R., Al Khatib, H., et al.: Human postprandial responses to food and potential for precision nutrition. Nature Medicine 26(6), 964–973 (2020)

[8] Müllner, E., Röhnisch, H.E., Von Brömssen, C., Moazzami, A.A.: Metabolomics analysis reveals altered metabolites in lean compared with obese adolescents and additional metabolic shifts associated with hyperinsulinaemia and insulin resistance in obese adolescents: A cross-sectional study. Metabolomics 17(1), 1–13 (2021)

[9] Vis, D.J., Westerhuis, J.A., Jacobs, D.M., Duynhoven, J.P., Wopereis, S., Ommen, B., Hendriks, M.M., Smilde, A.K.: Analyzing metabolomics-based challenge tests. Metabolomics 11(1), 50–63 (2015)

[10] Smilde, A.K., Jansen, J.J., Hoefsloot, H.C., Lamers, R.-J.A., Van Der Greef, J., Timmerman, M.E.: Anova-simultaneous component analysis (asca): a new tool for analyzing designed metabolomics data. Bioinformatics 21(13), 3043–3048 (2005)

[11] Harrington, P., Vieira, N.E., Espinoza, J., Nien, J.K., Romero, R., Yergey, A.L.: Analysis of variance–principal component analysis: A soft tool for proteomic discovery. Analytica Chimica Acta 544(1-2), 118–127 (2005)

[12] Thissen, U., Wopereis, S., Berg, S.A., Bobeldijk, I., Kleemann, R., Kooistra, T., Dijk, K., Ommen, B., Smilde, A.K.: Improving the analysis of designed studies by combining statistical modelling with study design information. BMC Bioinformatics 10(1), 1–15 (2009)

[13] Thiel, M., Feraud, B., Govaerts, B.: Asca+ and apca+: Extensions of asca and apca in the analysis of unbalanced multifactorial designs. Journal of Chemometrics 31(6), 2895 (2017)

[14] Martin, M., Govaerts, B.: Limm-pca: Combining asca+ and linear mixed models to analyse high-dimensional designed data. Journal of Chemometrics 34(6), 3232 (2020)

[15] Pellis, L., van Erk, M.J., van Ommen, B., Bakker, G.C.M., Hendriks, H.F.J., Cnubben, N.H.P., Kleemann, R., van Someren, E.P., Bobeldijk, I., Rubingh, C.M., Wopereis, S.: Plasma metabolomics and proteomics profiling after a postprandial challenge reveal subtle diet effects on human metabolic status. Metabolomics 8, 347–359 (2012)

[16] Kolda, T.G., Bader, B.W.: Tensor decompositions and applications. SIAM Review 51(3), 455–500 (2009)

[17] Acar, E., Yener, B.: Unsupervised multiway data analysis: A literature survey. IEEE Transactions on Knowledge and Data Engineering 21(1), 6–20 (2009)

[18] Smilde, A., Bro, R., Geladi, P.: Multi-Way Analysis: Applications in the Chemical Sciences. Wiley, West Sussex, England (2004)

[19] Harshman, R.A.: Foundations of the parafac procedure: Models and conditions for an “explanatory” multimodal factor analysis. UCLA working papers in phonetics 16, 1–84 (1970)

[20] Carroll, J.D., Chang, J.-J.: Analysis of individual differences in multidimensional scaling via an n-way generalization of “eckart-young” decomposition. Psychometrika 35(3), 283–319 (1970)

[21] Kruskal, J.B.: Three-way arrays: rank and uniqueness of trilinear decompositions, with application to arithmetic complexity and statistics. Linear Algebra and its Applications 18(2), 95–138 (1977)

[22] Bro, R.: PARAFAC. Tutorial and applications. Chemometrics and Intelligent Laboratory Systems 38(2), 149–171 (1997)

[23] Martino, C., Shenhav, L., Marotz, C.A., Armstrong, G., McDonald, D., Vázquez-Baeza, Y., Morton, J.T., Jiang, L., Dominguez-Bello, M.G., Swafford, A.D., et al.: Context-aware dimensionality reduction deconvolutes gut microbial community dynamics. Nature Biotechnology 39(2), 165–168 (2021)

[24] Gardlo, A., Smilde, A.K., Hron, K., Hrda, M., Karlikova, R., Friedeckỳ, D., Adam, T.: Normalization techniques for parafac modeling of urine metabolomic data. Metabolomics 12(12), 1–13 (2016)

[25] Li, L., Hoefsloot, H., Graaf, A.A., Acar, E., Smilde, A.K.: Exploring dynamic metabolomics data with multiway data analysis: a simulation study. BMC Bioinformatics 23(31), 1–22 (2022)

[26] Twisk, J., Bosman, L., Hoekstra, T., Rijnhart, J., Welten, M., Heymans, M.: Different ways to estimate treatment effects in randomised controlled trials. Contemporary Clinical Trials Communications 10, 80–85 (2018)

[27] Mattes, R.D.: Oral fat exposure alters postprandial lipid metabolism in humans. The American Journal of Clinical Nutrition 63(6), 911–917 (1996)

[28] Wopereis, S., Wolvers, D., Erk, M., Gribnau, M., Kremer, B., Dorsten, F.A., Boelsma, E., Garczarek, U., Cnubben, N., Frenken, L., et al.: Assessment of inflammatory resilience in healthy subjects using dietary lipid and glucose challenges. BMC Medical Genomics 6, 44 (2013)

[29] Kurata, H.: Virtual metabolic human dynamic model for pathological analysis and therapy design for diabetes. iScience 24(2), 102101 (2021)

[30] Bisgaard, H.: The Copenhagen Prospective Study on Asthma in Childhood (COP-SAC): design, rationale, and baseline data from a longitudinal birth cohort study. Annals of Allergy, Asthma & Immunology 93(4), 381–389 (2004)

[31] Stroeve, J.H., Wietmarschen, H., Kremer, B.H., Ommen, B., Wopereis, S.: Phenotypic flexibility as a measure of health: the optimal nutritional stress response test. Genes & Nutrition 10(3), 1–21 (2015)

[32] Hitchcock, F.L.: The expression of a tensor or a polyadic as a sum of products. Journal of Mathematics and Physics 6(1), 164–189 (1927)

[33] Papalexakis, E.E., Faloutsos, C., Sidiropoulos, N.D.: Tensors for data mining and data fusion: Models, applications, and scalable algorithms. ACM Trans. Intell. Syst. Technol. 8(2) (2016)

[34] Acar, E., Bingol, C.A., Bingol, H., Bro, R., Yener, B.: Multiway analysis of epilepsy tensors. Bioinformatics 23(13), 10–18 (2007)

[35] Williams, A.H., Kim, T.H., Wang, F., Vyas, S., Ryu, S.I., Shenoy, K.V., Schnitzer, M., Kolda, T.G., Ganguli, S.: Unsupervised discovery of demixed, low-dimensional neural dynamics across multiple timescales through tensor component analysis. Neuron 98(6), 1099–1115 (2018)

[36] Tomasi, G., Bro, R.: Parafac and missing values. Chemometrics and Intelligent Laboratory Systems 75(2), 163–180 (2005)

[37] Acar, E., Dunlavy, D.M., Kolda, T.G., Mørup, M.: Scalable tensor factorizations for incomplete data. Chemometrics and Intelligent Laboratory Systems 106(1), 41–56 (2011)

[38] Håastad, J.: Tensor rank is np-complete. Journal of Algorithms 11(4), 644–654 (1990)

[39] Adali, T., Kantar, F., Akhonda, M.A.B.S., Strother, S., Calhoun, V.D., Acar, E.: Reproducibility in matrix and tensor decompositions: Focus on model match, interpretability, and uniqueness. IEEE Signal Processing Magazine 39(4), 8–24 (2022)

[40] Bro, R., Kiers, H.A.: A new efficient method for determining the number of components in parafac models. Journal of Chemometrics 17(5), 274–286 (2003)

[41] Harshman, R.A., De Sarbo, W.S.: An application of PARAFAC to a small sample problem, demonstrating preprocessing, orthogonality constraints, and split-half diagnostic techniques in Research Methods for Multimode Data Analysis. Praeger: New York, 602–642 (1984)

[42] Bro, R., Smilde, A.K.: Centering and scaling in component analysis. Journal of Chemometrics 17(1), 16–33 (2003)

[43] Acar, E., Dunlavy, D.M., Kolda, T.G.: A scalable optimization approach for fitting canonical tensor decompositions. Journal of Chemometrics 25(2), 67–86 (2011)

[44] Bader, B.W., Kolda, T.G., et al.: Matlab Tensor Toolbox, Version 3.1. https://www.tensortoolbox.org

[45] Kahler, A., Zimmermann, M., Langhans, W.: Suppression of hepatic fatty acid oxidation and food intake in men. Nutrition 15(11-12), 819–828 (1999)

[46] Saito, K., Maekawa, K., Pappan, K.L., Urata, M., Ishikawa, M., Kumagai, Y., Saito, Y.: Differences in metabolite profiles between blood matrices, ages, and sexes among caucasian individuals and their inter-individual variations. Metabolomics 10(3), 402–413 (2014)

[47] Smilde, A.K., Werf, M.J., Schaller, J.-P., Kistemaker, C.: Characterizing the precision of mass-spectrometry-based metabolic profiling platforms. Analyst 134(11), 2281–2285 (2009)

[48] Vilozni, D., Bentur, L., Efrati, O., Barak, A., Szeinberg, A., Shoseyov, D., Yahav, Y., Augarten, A.: Exercise challenge test in 3-to 6-year-old asthmatic children. Chest 132(2), 497–503 (2007)

[49] Harshman, R.A.: Parafac2: Mathematical and technical notes. UCLA working papers in phonetics 22, 30–44 (1972)

[50] Madsen, K.H., Churchill, N.W., Mørup, M.: Quantifying functional connectivity in multi-subject fMRI data using component models. Human Brain Mapping 38, 882–899 (2017)

[51] Roald, M., Bhinge, S., Jia, C., Calhoun, V., Adali, T., Acar, E.: Tracing network evolution using the PARAFAC2 model. In: ICASSP’20, pp. 1100–1104 (2020)

[52] Acar, E., Roald, M., Hossain, K.M., Calhoun, V.D., Adali, T.: Tracing evolving networks using tensor factorizations vs. ica-based approaches. Frontiers in Neuroscience 16, 861402 (2022)

[53] Bro, R., Andersson, C.A., Kiers, H.A.: Parafac2—part ii. modeling chromatographic data with retention time shifts. Journal of Chemometrics 13, 295–309 (1999)

[54] Jendoubi, T., Ebbels, T.M.D.: Integrative analysis of time course metabolic data and biomarker discovery. BMC Bioinformatics 21, 11 (2020)

