## Additional file 1 for "Analyzing postprandial metabolomics data using multiway models: A simulation study"

**A list of parameters that are randomly perturbated for generating different individuals**

Vdif_glut2_Glc_B_L
Vmax_gk_Glc_L
Vmax_g6pase_G6p_L
Vmax_ppp_G6p_L
Vmax_gs_G6p_L
Vmax_gd_Glygn_L
Vmax_pfk_G6p_L
Vmax_fbp_Gap_L
Vmax_pk_Gap_L
Vmax_pepck_Pyr_L
Vdif_pyrt_Pyr_B_L
Vdif_lact_Lac_B_L
Vmax_ldh_Pyr_L
Vdif_alat_Ala_B_L
Vmax_alata_Pyr_L
Vmax_pdh_Pyr_L
Vmax_tca_Accoa_LM
Vdif_ffat_FFA_B_L
Vmax_ffat_FFA_B_L
Vmax_tgsyn_FFA_L
Vmax_tgdeg_TG_L
Vmax_glyk_Glyc_L
Vdif_tgt_TG_B_L
Vmax_tgt_TG_B_L
Vmax_g3pd_Gap_L
Vmax_boxid_FFA_L
Vdif_glyct_Glyc_B_L
Vmax_accoat_Accoa_LM_LC
Vmax_bhbsyn_Accoa_LM
Vdif_bhbt_Bhb_B_L
Vmax_lipog1_Accoa_LC
Vmax_lipog2_Malcoa_L
Vmax_cholsyn1_Accoa_LC
Vmax_cholsyn2_Hmgcoa_L
Vmax_cholt_Chol_L
Vmax_atpsynf_Fadh_L
Vmax_atpsynn_Nadh_L
Vmax_nadhk_Nadh_L
Vmax_atpuse_Atp_L
Vmax_utpuse_Utp_L
Vmax_gtpuse_Gtp_L
Vmax_nadhuse_Nadh_L
Vmax_nadphuse_Nadph_L
Vmax_fadhuse_Fadh_L
Vmax_ampreg_Amp_L
Vmax_gdpreg_Gdp_L
Vmax_udpreg_Udp_L
Vmax_ck_Cre_L
