## Additional file 2 for "Analyzing postprandial metabolomics data using multiway models: A simulation study"

### Supplementary figures and tables for the main manuscript

#### Supplemental figures

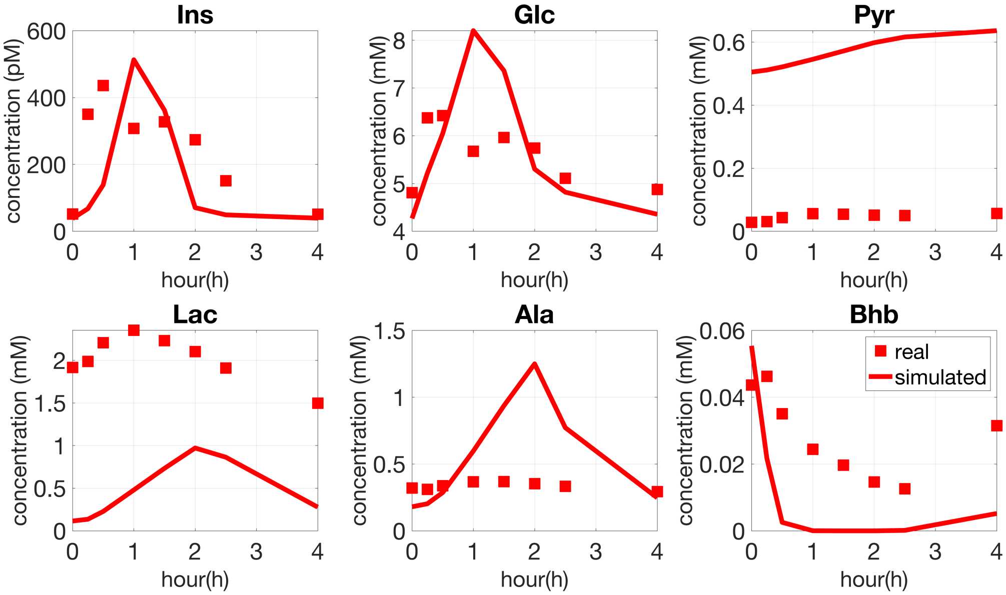


Figure S.1: **Median time profiles of the real vs. simulated (50 control subjects and with the random perturbation level for individual variation set to $\alpha = 0.2$) data generated from the default human whole-body metabolic model.**

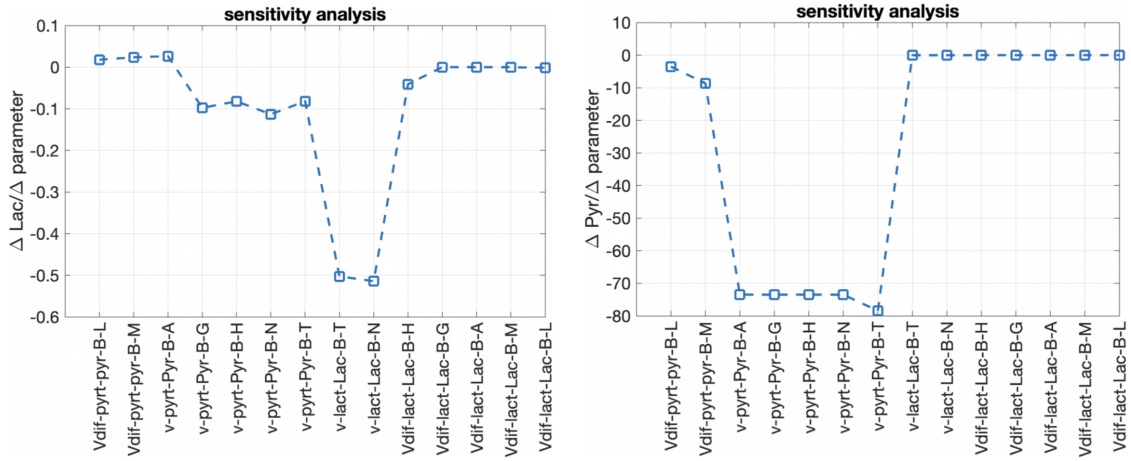


Figure S.2: **Sensitivity analysis for metabolites Lac (left panel) and Pyr (right panel). We observed an apparent deviation of Pyr from the reference range** $[0.04, 0.1]$ **mmol/L using the default model, as shown in the figure in Additional file 1; therefore, we performed a sensitivity analysis (checking how the change of parameters related to Pyr will affect the change of its 10h-fasting concentration) to tune the parameters in the human whole-body model so that the 10h-fasting concentration of Pyr falls into the reference range. From Figure S.2 (right panel), we select to increase v-pyrt-pyr-B-A by 0.007 (the default value is 0) to obtain a significant decrease of the 10h-fasting Pyr. In addition, we increase Vdif-pyrt-pyr-B-L by 0.003 (the default value is 0) to make an adjustment. Although the left panel in S.2 indicates decreasing v-lact-lac-B-N or v-lact-lac-B-T will increase the 10h-fasting Lac, such an increase will lead to negative concentrations of Lactate in the brain (N) or other tissues (T). We make a minor adjustment to Lac by setting Vdif-lact-Lac-B-L to 12*0.08 (the default value is 12). Details about the equations and parameters related to Pyr and Lac can be found in [1].**

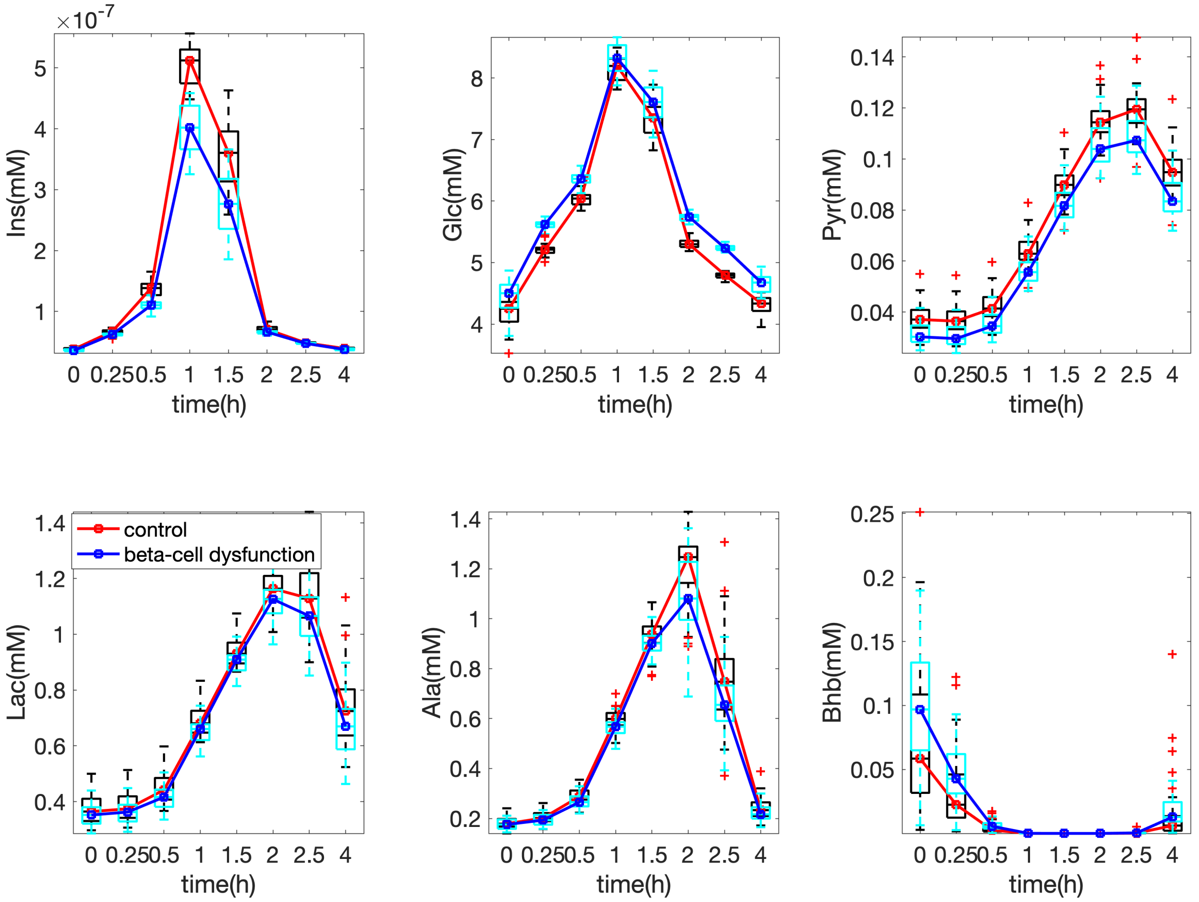


Figure S.3: **Median time profiles of 50 control vs. 50 *beta-cell dysfunction* subjects for the *full-dynamic* data with individual variations introduced by setting the random perturbation level to $\alpha = 0.2$.**

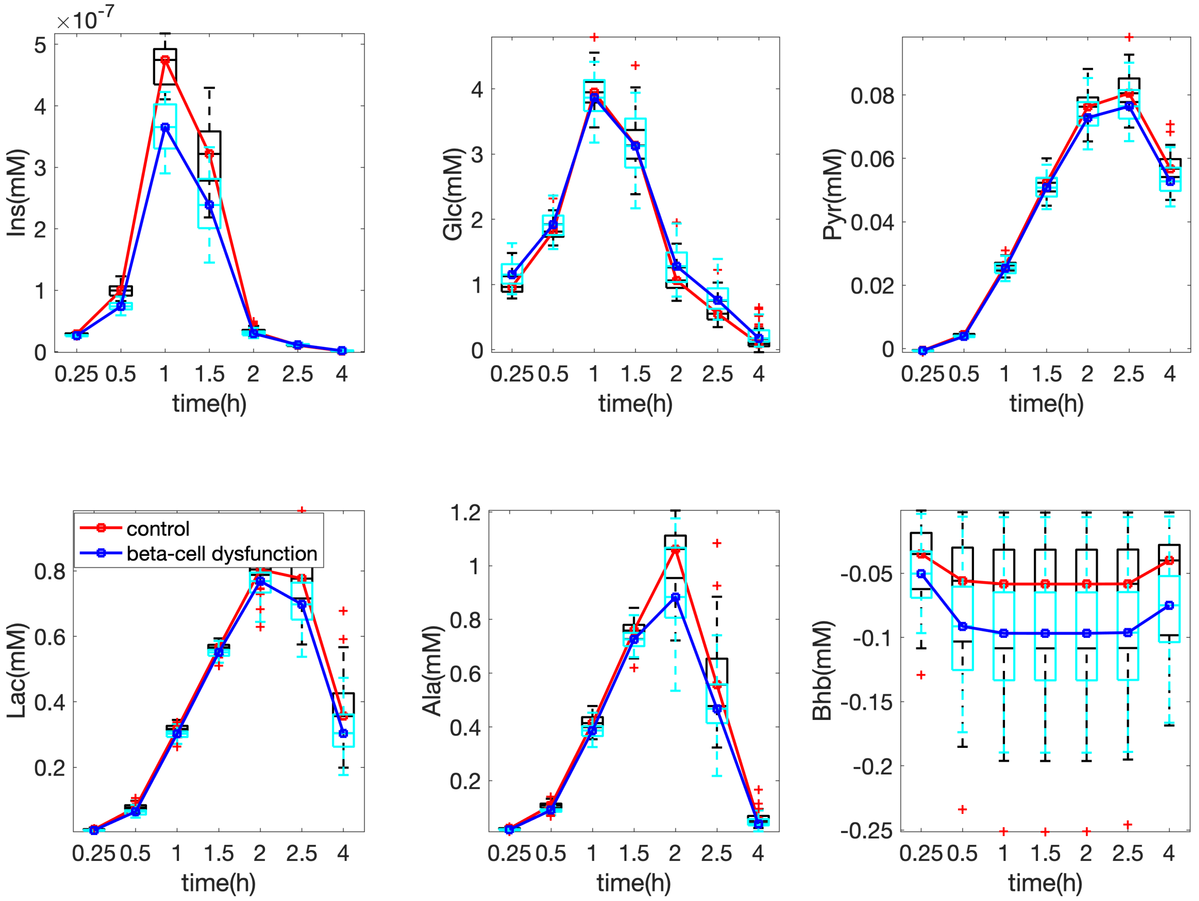


Figure S.4: **Median time profiles of 50 control vs. 50 *beta-cell dysfunction* subjects for the *T0-corrected* data with individual variations introduced by setting the random perturbation level to $\alpha = 0.2$.**

### Supplemental Tables

| | glucose (g) | fat (g) | protein (g) |
|---|---|---|---|
| real data | 60 | 75 | 25 |
| simulated data | 87 | 33 | 0 |

Table S.1: **Meal compositions of the real and simulated data.**

| | **Individual Variation** | | **R** | **FMS** |
|---|---|---|---|---|
| *insulin resistance* | $\alpha = 0.2$ | balanced | 4 | - |
| | | unbalanced | 4 | 1.00 |
| | $\alpha = 0.4$ | balanced | 4 | 0.97 |
| | | unbalanced | 4 | 0.97 |
| *beta-cell dysfunction* | $\alpha = 0.2$ | balanced | 4 | - |
| | | unbalanced | 4 | 1.00 |
| | $\alpha = 0.4$ | balanced | 5 | 0.93 |
| | | unbalanced | 5 | 0.90 |

Table S.2: ***Full-dynamic data analysis.*** **Similarity between the CP factors (in the *metabolites* and *time* modes) extracted from different data sets and the CP factors of the data set with $\alpha = 0.2$ and balanced samples. Here, $\alpha$ denotes the level of individual variation, where a smaller number indicates a lower level of individual variation.**

| | **Individual Variation** | | **R** | **FMS** |
|---|---|---|---|---|
| *insulin resistance* | $\alpha = 0.2$ | balanced | 4 | - |
| | | unbalanced | 4 | 1.00 |
| | $\alpha = 0.4$ | balanced | 4 | 0.99 |
| | | unbalanced | 4 | 0.99 |
| *beta-cell dysfunction* | $\alpha = 0.2$ | balanced | 4 | - |
| | | unbalanced | 4 | 0.96 |
| | $\alpha = 0.4$ | balanced | 5 | 0.95 |
| | | unbalanced | 5 | 0.95 |

Table S.3: ***T0-corrected data analysis.*** **Similarity between the CP factors (in the *metabolites* and *time* modes) extracted from different data sets and the CP factors of the data set with $\alpha = 0.2$ and balanced samples. Here, $\alpha$ denotes the level of individual variation, where a smaller number indicates a lower level of individual variation.**

### References

[1] Hiroyuki Kurata. Virtual metabolic human dynamic model for pathological analysis and therapy design for diabetes. *iScience*, 24(2):102101, 2021.
