## Additional file 3 for "Analyzing postprandial metabolomics data using multiway models: A simulation study"

### Selection of the number of components in CP models

When selecting the number of components (i.e., $R$) in CP models, we check the increase in model fit, drop in core consistency, and replicability of the model, which are discussed in the following. In addition, the selection also depends on the interpretability of the model.

#### Model fit

The *model fit* is defined as follows:

$$\text{Fit (\%)} = (1 - \frac{\left\| \mathscr{X} - \hat{\mathscr{X}} \right\|^2}{\| \mathscr{X} \|^2}) \times 100,$$

where $\mathscr{X}$ and $\hat{\mathscr{X}}$ correspond to the data tensor and approximation of the data by the model, respectively. A fit value close to 100% means that data $\mathscr{X}$ is well explained by the model; otherwise, there is an unexplained part left in the residuals. If a significant gain in model fit is observed as the number of components increases, we should consider having more components.

#### Core consistency

The core consistency diagnostic [1] is another approach for determining the number of components in a CP model. The core consistency compares the core array of the CP model with the core array obtained by modeling the data using a Tucker3 model [2] based on the CP factors. A core consistency value close to 100% indicates an appropriate number of components, while too many components may lead to a drop in the core consistency value.

#### Replicability

Another approach to selecting the number of components in a CP model is to check whether the model produces identical patterns from independent subsamples of the data set, e.g., as in split-half analysis [3]. Here, we propose a method to check the replicability of a model, which can be viewed as an extension of split-half analysis. The replicability is tested by leaving out random subsets of subjects (randomly selecting and leaving out 10% of the total subjects) and computing the similarity of the factors in the *metabolites* and *time* modes extracted using a CP model from the remaining data. We quantify the similarity of CP models from two splits using the factor match score (FMS) introduced in the main text. The model is considered replicable if the patterns are similar enough. Figure A.1 illustrates the proposed replicability check.

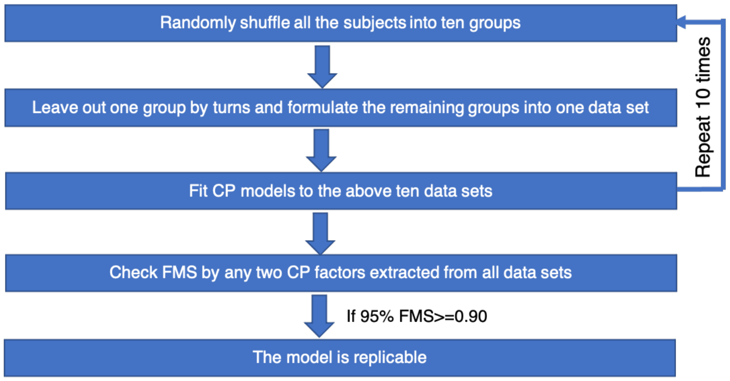


Figure A.1: **The replicability test used to select the number of components.**

#### Model selection for each data set

##### ***Full-dynamic*** analysis for the data with ***insulin-resistant*** vs. control group, $\alpha = 0.2$, balanced samples

The model fit increases evidently from $R = 1$ to $R = 3$ (see Figure A.2a). The core consistency drops significantly when we increase the number of components from $R = 5$ to $R = 6$ (see Figure A.2b).In addition, the 6-component model can not be replicated (see Figure A.3f). Therefore, we select among models with $R \leq 5$. We choose the 4-component model since it is more interpretable and can be replicated (see Figure A.3d). Compared with the 3-component model, the 4-component model captures an extra constant temporal pattern (i.e., the fourth component), which reflects the fasting-state information (see the comparison of Figure A.4 and A.5). The 5-component model splits one of the patterns in the 4-component model, i.e., the first component ($\mathbf{a}_1$, $\mathbf{b}_1$ and $\mathbf{c}_1$ in Figure A.5), where Lac and Ala have large coefficients on $\mathbf{b}_1$, into two components, i.e., the first and fifth components in the 5-component model (Figure A.6).

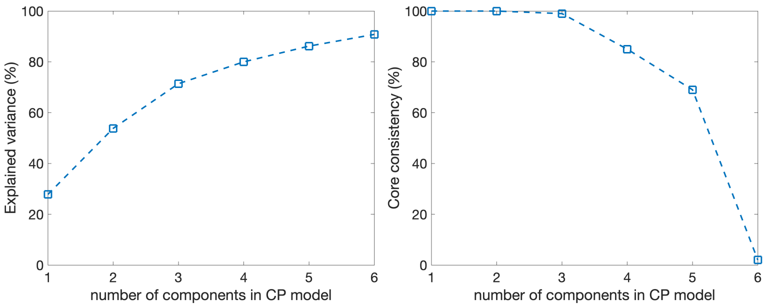


(a) (b)

Figure A.2: **(a) Model fit, and (b) core consistency of CP models using different numbers of components for the *full-dynamic* data with *insulin-resistant* vs. control group, $\alpha = 0.2$ and balanced samples.**

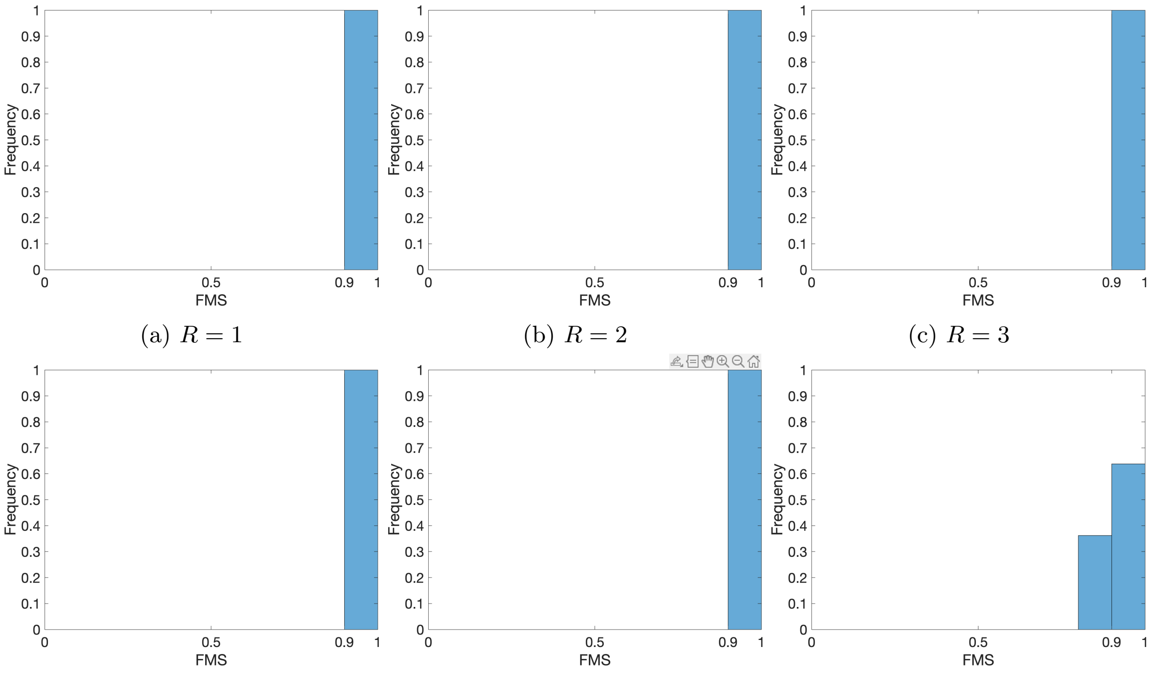


(a) $R = 1$ (b) $R = 2$ (c) $R = 3$

(d) $R = 4$ (e) $R = 5$ (f) $R = 6$

Figure A.3: **Histogram of FMS values between CP factors (in the *metabolites* and *time* modes) extracted from all splits of the *full-dynamic* data with *insulin-resistant* vs. control group, $\alpha = 0.2$ and balanced samples.**

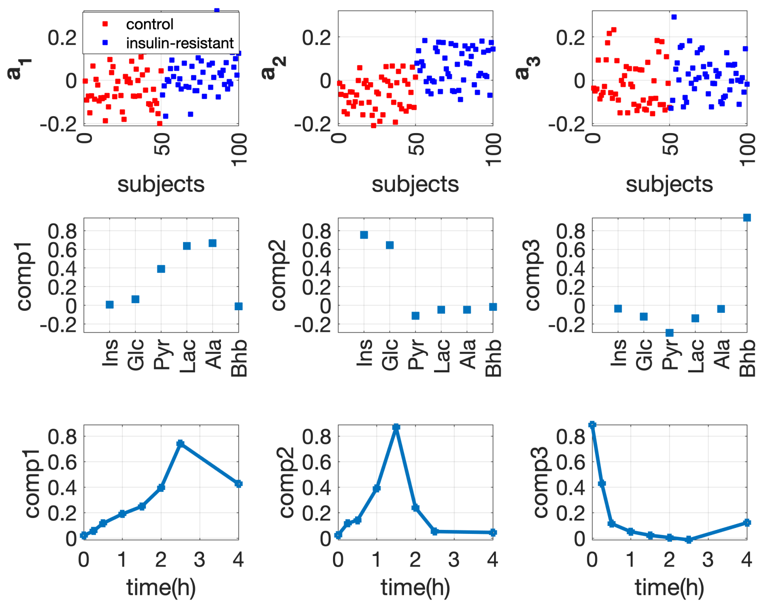


Figure A.4: **Factors of the 3-component CP model for the *full-dynamic* data with *insulin-resistant* vs. control group and $\alpha = 0.2$.**

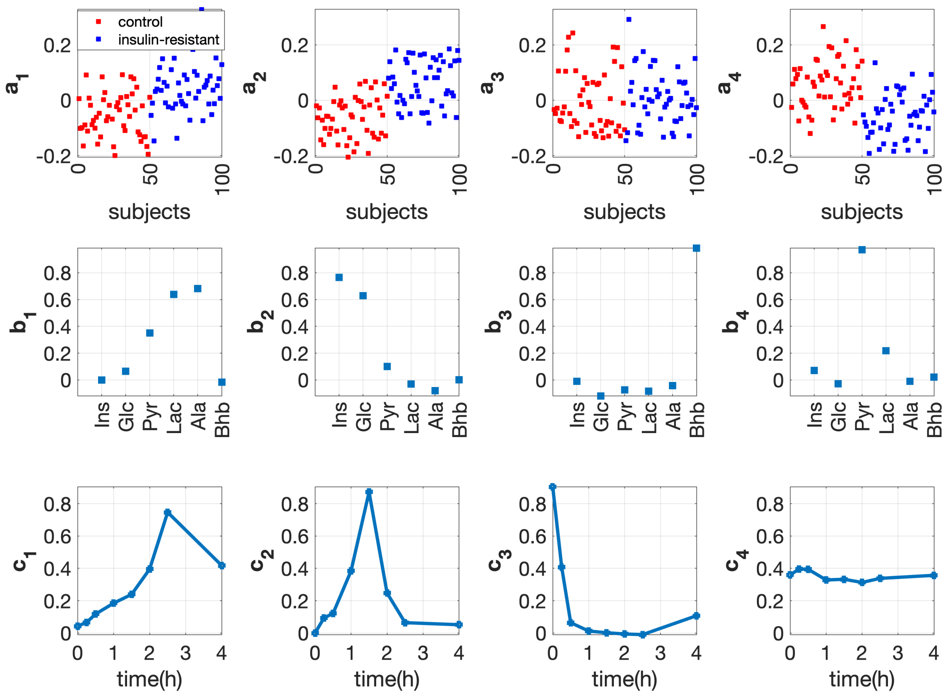


Figure A.5: **Factors of the 4-component CP model for the *full-dynamic* data with *insulin-resistant* vs. control group and $\alpha = 0.2$.**

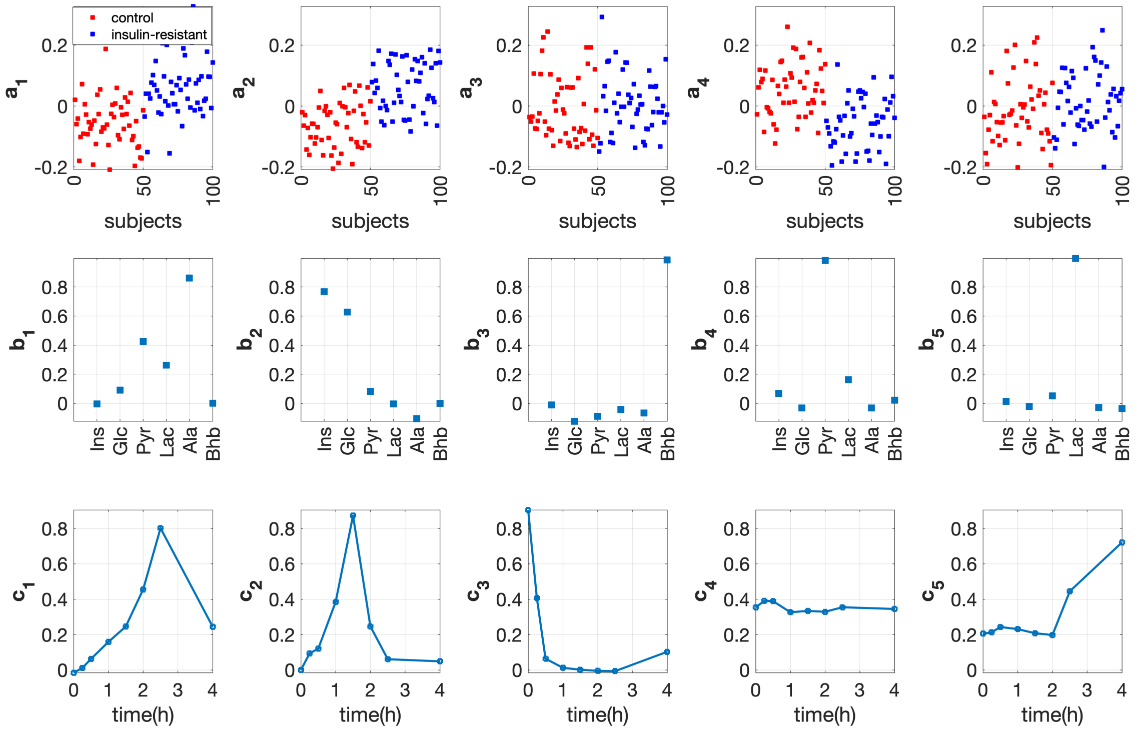


Figure A.6: **Factors of the 5-component CP model for the *full-dynamic* data with *insulin-resistant* vs. control group and $\alpha = 0.2$.**

**T0-corrected analysis for the data with *insulin-resistant* vs. control group, $\alpha = 0.2$ and balanced samples**

The model fit increases evidently from $R = 1$ to $R = 3$ (see Figure A.7a). The core consistency drops evidently when we increase the number of components from $R = 5$ to $R = 6$ (see Figure A.7b); therefore, we consider models with $R \leq 5$. The 5-component model cannot be replicated but the 4-component model can (see Figure A.8). Therefore, we may prefer the 4-component model. In addition, compared with the 3-component model, the 4-component model captures an extra dynamic pattern related to metabolite Pyr (see the comparison of Figure A.9 with A.10), which provides useful information in terms of understanding metabolic differences at the *fasting* vs. *pure-dynamic* state.

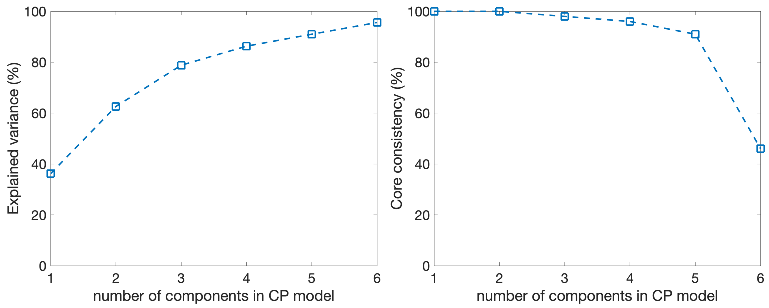


(a) (b)

Figure A.7: **(a) Model fit, and (b) core consistency of the CP models using different numbers of components for *T0-corrected* data generated with *insulin-resistant* vs. control group, $\alpha = 0.2$ and balanced samples.**

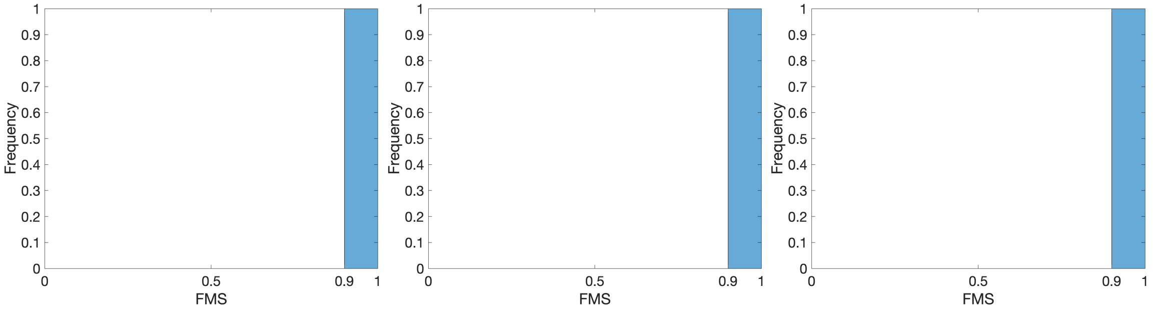


(a) $R = 1$ (b) $R = 2$ (c) $R = 3$

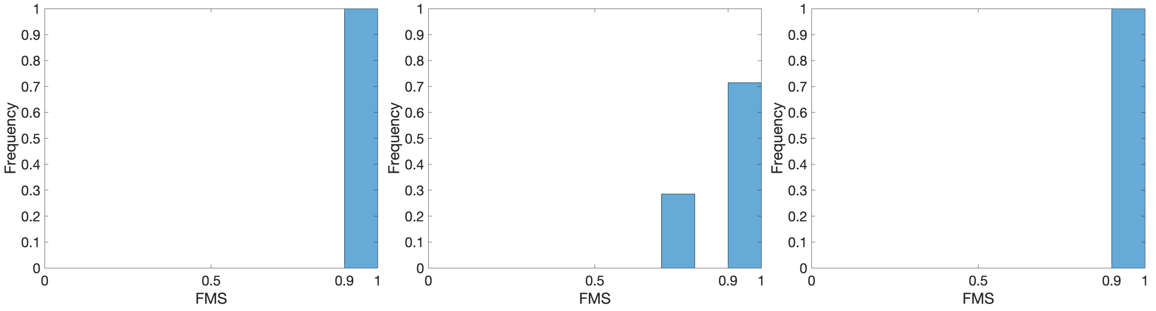


(d) $R = 4$ (e) $R = 5$ (f) $R = 6$

Figure A.8: **Histogram of FMS values between CP factors (in the *metabolites* and *time* modes) extracted from all splits of *T0-corrected* data with *insulin-resistant* vs. control group, $\alpha = 0.2$ and balanced samples.**

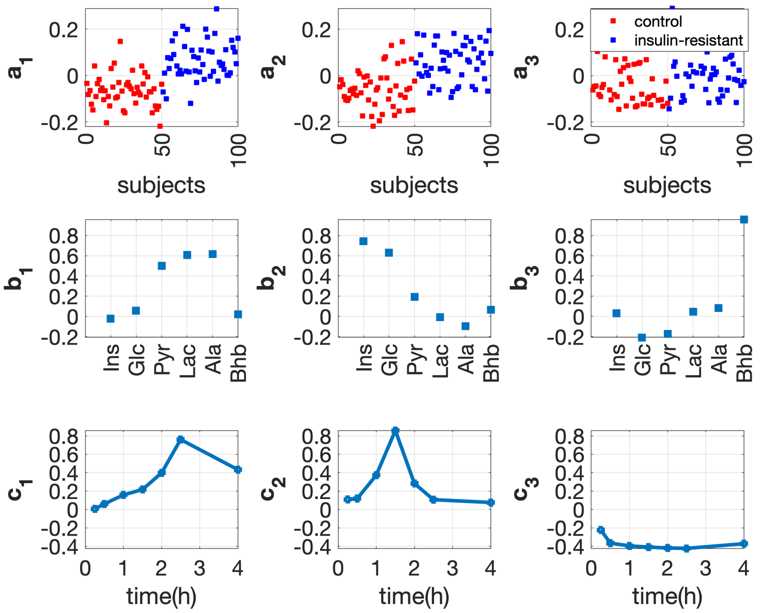


Figure A.9: **Factors of the 3-component CP model for the *T0-corrected* data with *insulin-resistant* vs. control group and $\alpha = 0.2$.**

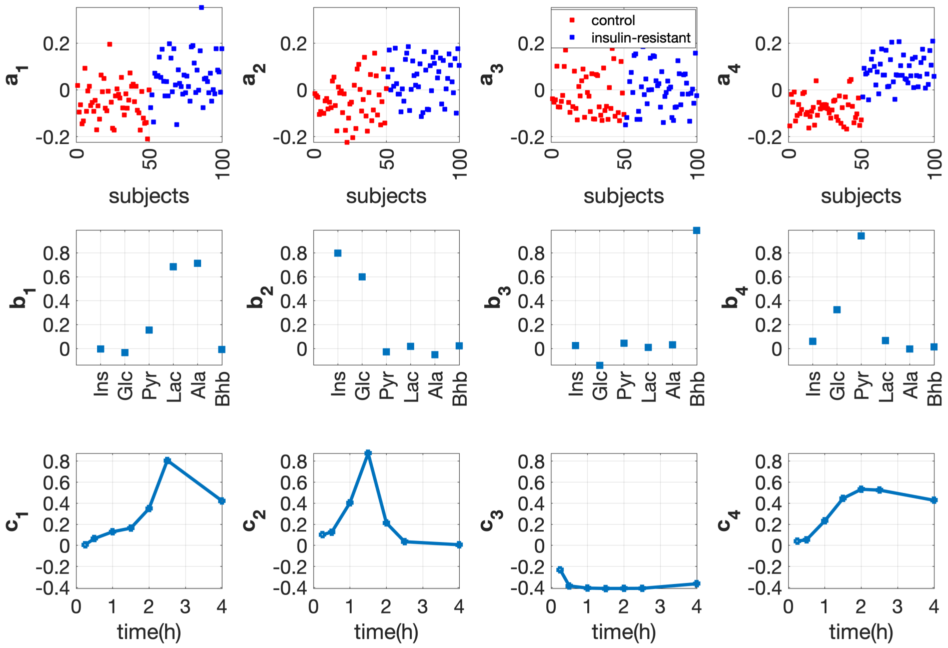


Figure A.10: **Factors of the 4-component CP model for the *T0-corrected* data with *insulin-resistant* vs. control group and $\alpha = 0.2$.**

**Full-dynamic analysis for the data generated with beta-cell dysfunction vs. control group, $\alpha = 0.2$ and balanced samples**

The model fit increases evidently from $R = 1$ to $R = 4$ (see Figure A.11a). The core consistency drops clearly when we increase the number of components from $R = 4$ to $R = 5$ and from $R = 5$ to $R = 6$ (see Figure A.11b). This indicates considering CP models with $R = 4$ or $R = 5$. The 5-component model cannot be replicated but the 4-component model can (see Figure A.12). Therefore, we select the 4-component model. In addition, compared with the 3-component model, the 4-component model captures the subject group separation much better (see Figure A.13 vs. A.14).

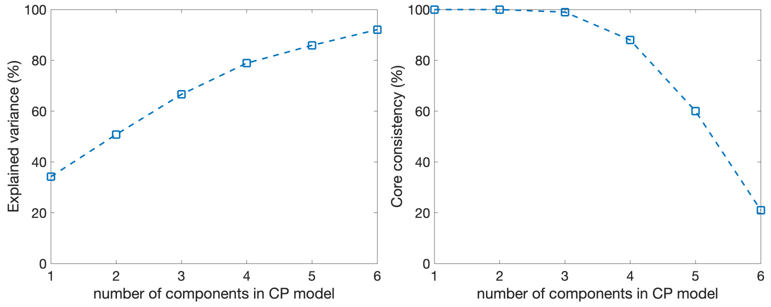


(a) (b)

Figure A.11: **(a) Model fit, and (b) core consistency of the CP models using different numbers of components for the (*full-dynamic*) data with *beta-cell dysfunction* vs. control group, $\alpha = 0.2$ and balanced samples.**

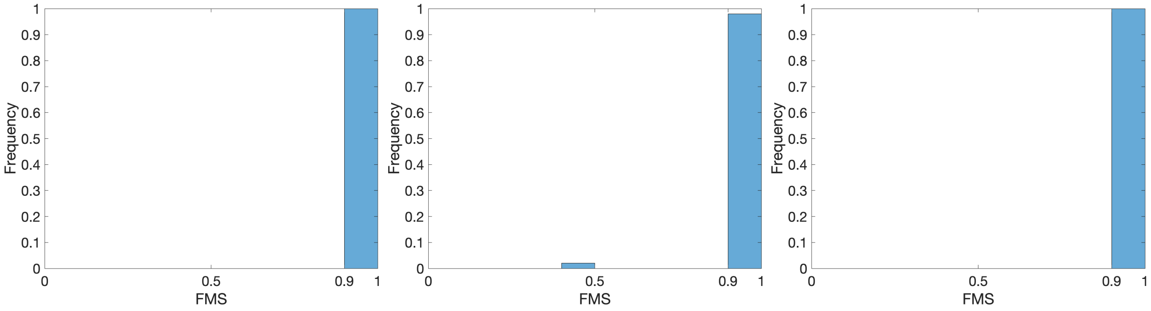


(a) 1-component CP model (b) 2-component CP model (c) 3-component CP model

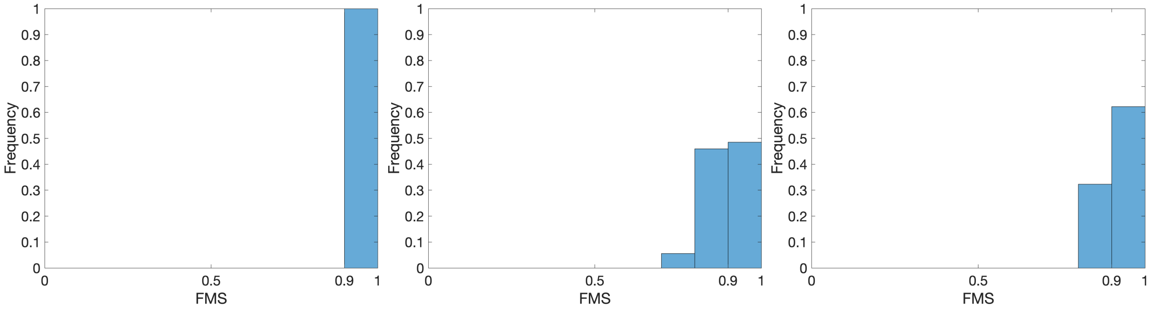


(d) $R = 4$ (e) $R = 5$ (f) $R = 6$

Figure A.12: **Histogram of FMS values between CP factors (in the *metabolites* and *time* modes) extracted from all splits of *full-dynamic* data with *beta-cell dysfunction* vs. control group, $\alpha = 0.2$ and balanced samples.**

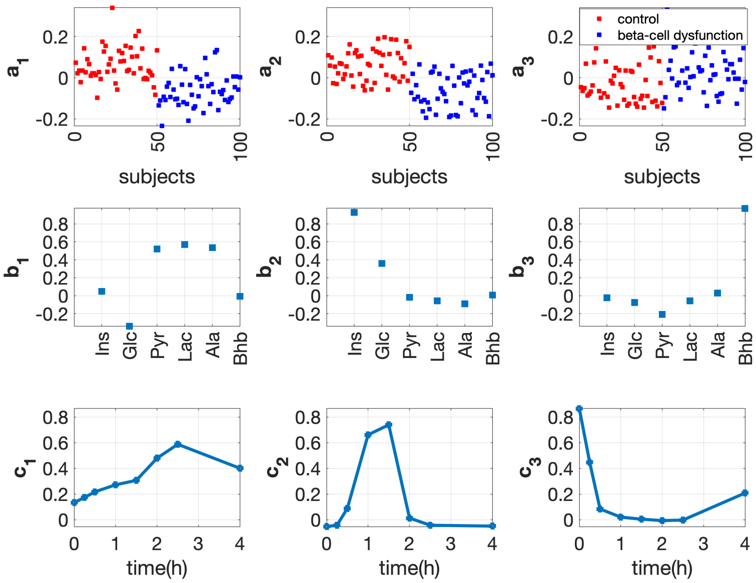


Figure A.13: **Factors of the 3-component CP model for the *full-dynamic* data with *beta-cell dysfunction* vs. control group, $\alpha = 0.2$ and balanced samples.**

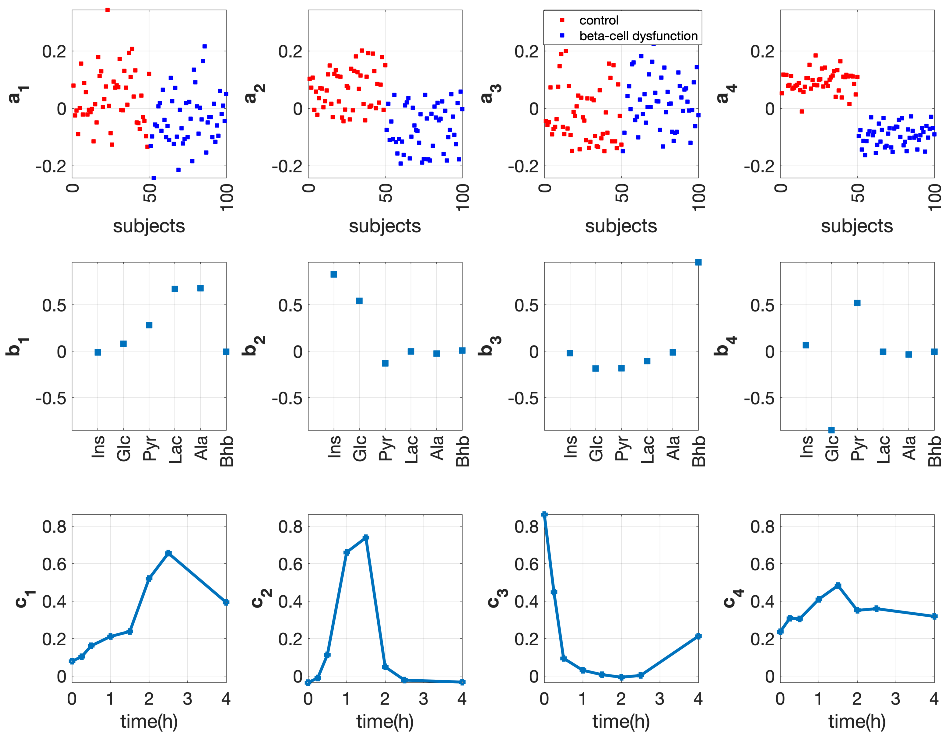


Figure A.14: **Factors of the 4-component CP model for the *full-dynamic* data with *beta-cell dysfunction* vs. control group, $\alpha = 0.2$ and balanced samples.**

##### *T0-corrected* analysis for the data with *beta-cell dysfunction* vs. control group, $\alpha = 0.2$ and balanced samples

The model fit increases evidently from $R = 1$ to $R = 3$ (see Figure A.15a). The core consistency drops clearly when we increase the number of components from $R = 3$ to $R = 4$ and from $R = 5$ to $R = 6$ (see Figure A.15b). Therefore, we consider models with $R = 3$, $R = 4$, or $R = 5$. The replicability check shows that both the 3-component and 5-component models can be reproduced well (see Figure A.16c and A.16e). For the 4-component model, a few splits have different patterns from the remaining splits, as illustrated in Figure A.16d. However, the 4-component model is more interpretable than the 3-component and 5-component models. Compared with the 3-component model, the 4-component model extracts Bhb out from the second component ($\mathbf{b}_2$) in the 3-component model in Figure A.17, which indicates that Bhb does not contribute to the subject group separation. The 5-component model splits the fourth component ($\mathbf{a}_4$, $\mathbf{b}_4$ and $\mathbf{c}_4$) in the 4-component model into two components (the fourth and fifth components in the 5-component model), which does not reveal that Glc and Pyr are close to each other (see Figure A.18 vs. A.19). Therefore, we prefer the 4-component model for this data set.

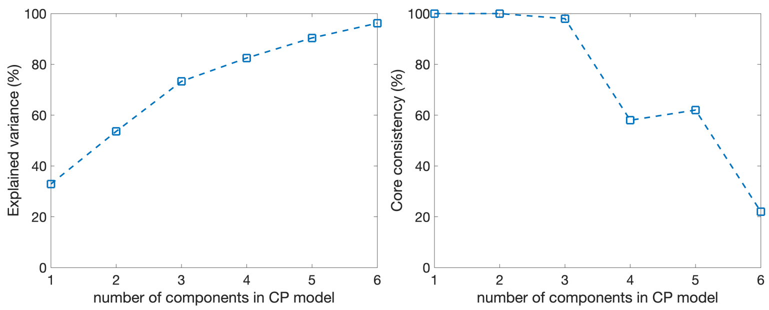


(a) (b)

Figure A.15: **(a) Model fit, and (b) core consistency of the CP models using different numbers of components for the *T0-corrected* data with *beta-cell dysfunction* vs. control group, $\alpha = 0.2$ and balanced samples**

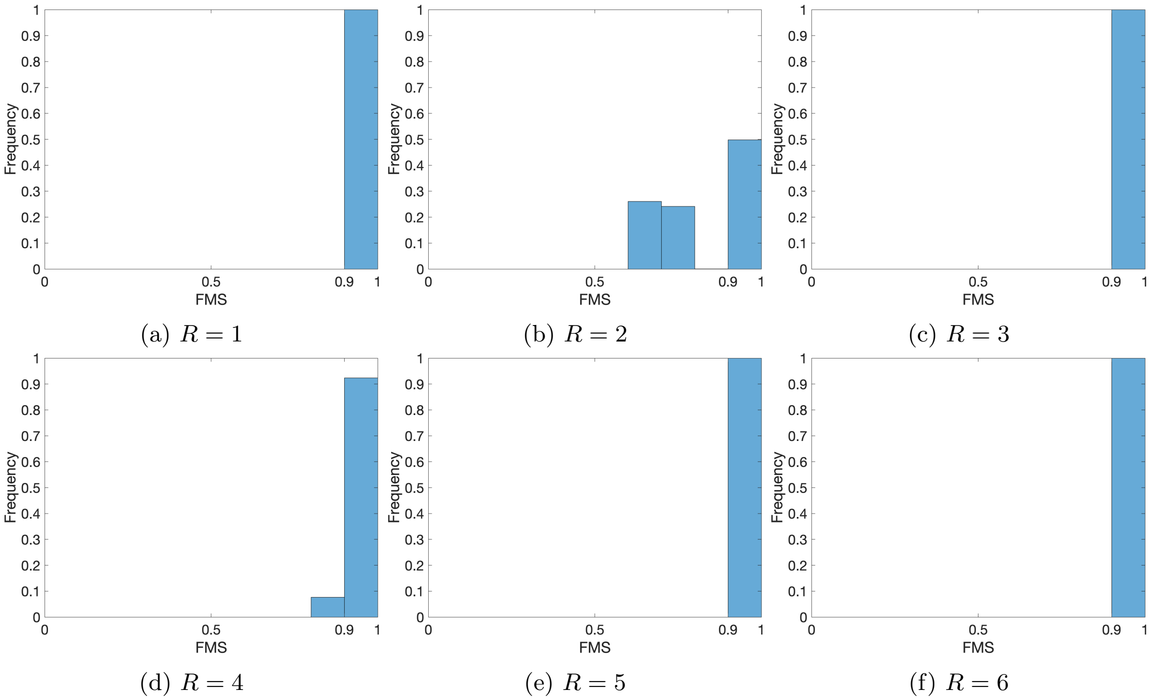


(a) $R = 1$ (b) $R = 2$ (c) $R = 3$

(d) $R = 4$ (e) $R = 5$ (f) $R = 6$

Figure A.16: **Histogram of FMS values between CP factors (in the *metabolites* and *time* modes) extracted from all splits of the *T0-corrected* data with *beta-cell dysfunction* vs. control group, $\alpha = 0.2$ and balanced samples.**

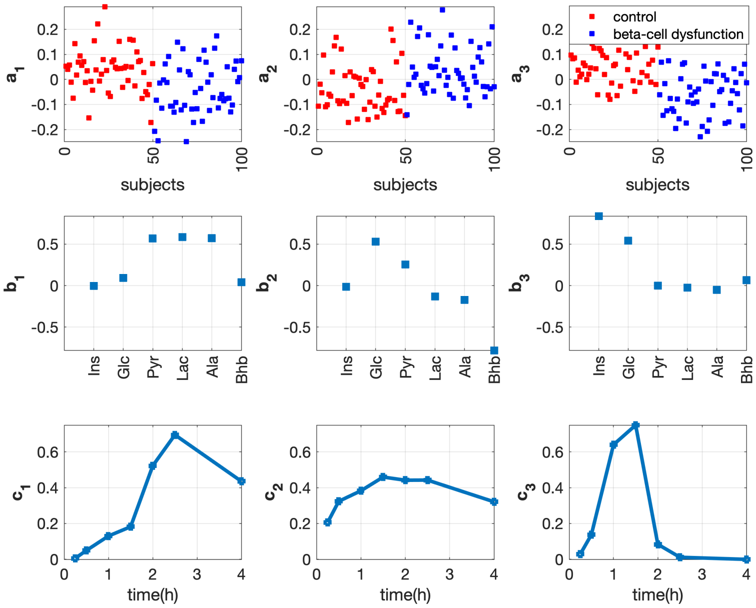


Figure A.17: **Factors of the 3-component CP model for the *T0-corrected* data with *beta-cell dysfunction* vs. control group and $\alpha = 0.2$.**

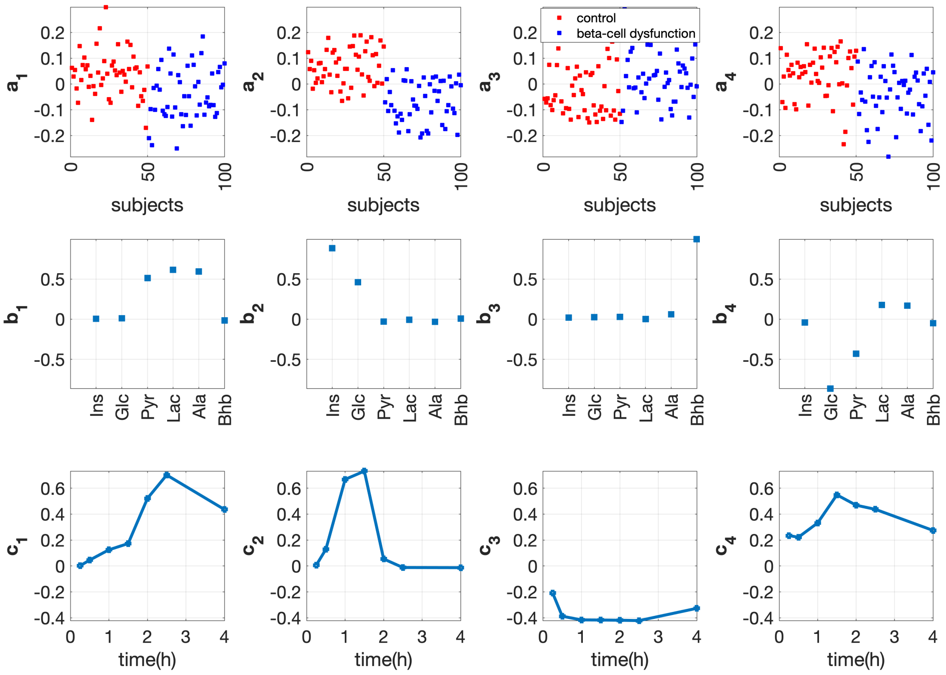


Figure A.18: **Factors of the 4-component CP model for the *T0-corrected* data with *beta-cell dysfunction* vs. control group and $\alpha = 0.2$.**

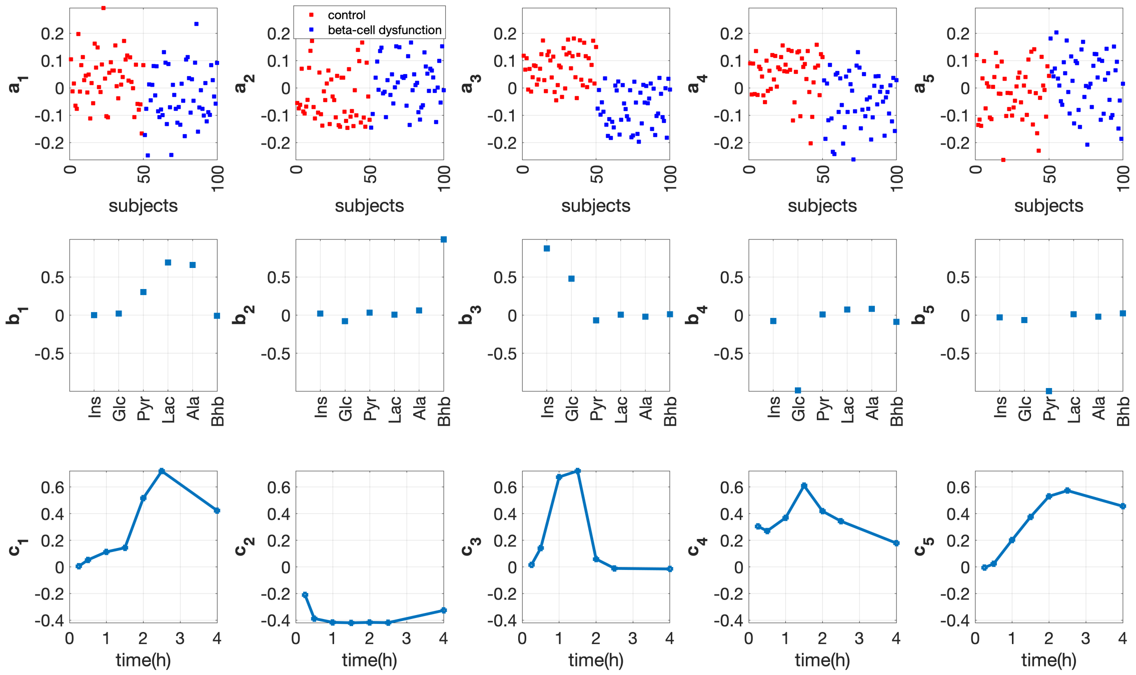


Figure A.19: **Factors of the 5-component CP model for the *T0-corrected* data with *beta-cell dysfunction* vs. control group and $\alpha = 0.2$.**

##### ***Full-dynamic* analysis for the real data consisting of 299 subjects and 6 measurements**

The model fit increases evidently from $R = 1$ to $R = 3$ (Figure A.20a). The core consistency drops significantly when we increase the number of components from $R = 3$ to $R = 4$ and from $R = 4$ to $R = 5$ (Figure A.20b). This indicates considering the models with $R = 3$ or $R = 4$. The 3-component model can be well replicated, and there are some unlucky splits when using the 4-component model (see Figure A.21). The main differences between the 4-component model from the 3-component model are Ala splits out from the third component in the 3-component model, and Pyr splits from the first component in the 3-component model to the third component in the 4-component model (see Figure A.22 vs. A.23). However, it makes sense that Pyr, Lac and Ala stay close since they are tied to each other with reactions Pyr↔Lac and Pyr↔Ala as shown in the pathway in the main text. Therefore, we prefer the 3-component CP model for this analysis.

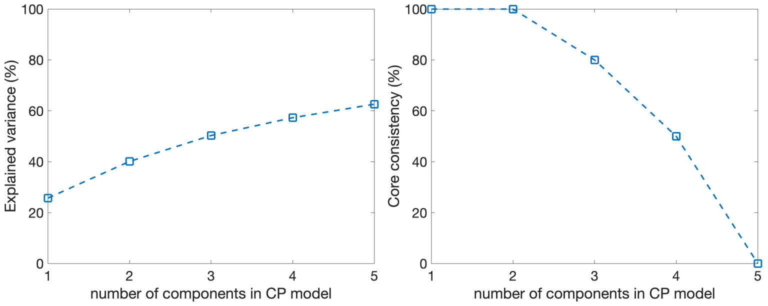


(a) (b)

Figure A.20: **(a) Model fit, and (b) core consistency of the CP models using different numbers of components for the real (*full-dynamic*) data set with 299 subjects and 6 metabolites.**

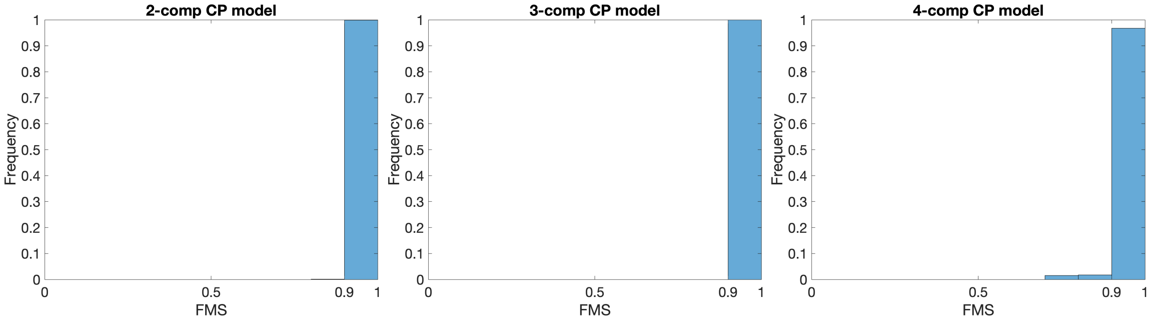


(a) $R = 2$ (b) $R = 3$ (c) $R = 4$

Figure A.21: **Histogram of FMS values between CP factors (in the *metabolites* and *time* modes) extracted from all splits of the real *full-dynamic* data consisting of 6 measurements.**

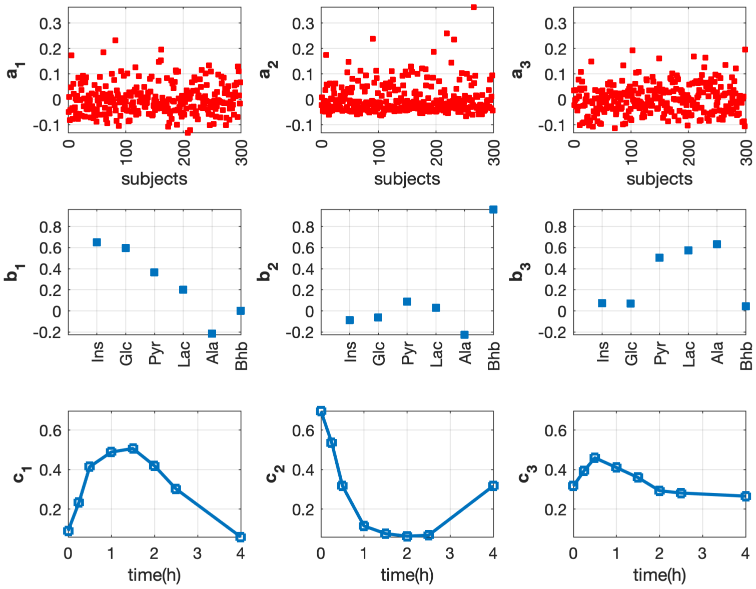


Figure A.22: **Factors of the 3-component CP model for the real data set with 6 measurements.**

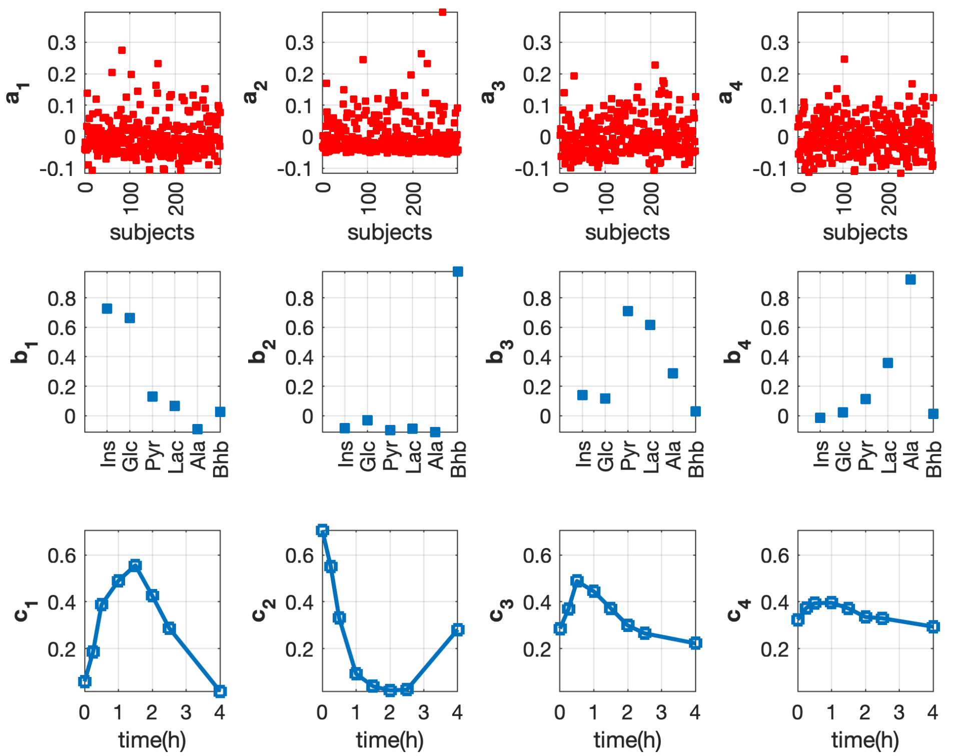


Figure A.23: **Factors of the 4-component CP model for the real data set with 6 measurements.**

### References


[1] Rasmus Bro and Henk AL Kiers. A new efficient method for determining the number of components in parafac models. *Journal of Chemometrics*, 17(5):274–286, 2003.

[2] Ledyard R Tucker. Some mathematical notes on three-mode factor analysis. *Psychometrika*, 31(3):279–311, 1966.

[3] R. A. Harshman and W. S. De Sarbo. An application of PARAFAC to a small sample problem, demonstrating preprocessing, orthogonality constraints, and split-half diagnostic techniques in *Research Methods for Multimode Data Analysis. Praeger: New York*, pages 602–642, 1984.
