## Additional file 4 for "Analyzing postprandial metabolomics data using multiway models: A simulation study"

### Comparison of CP models for real vs. simulated data

Figure 4 in the main text shows that patterns in the *metabolites* and *time* modes extracted by the 3-component CP model for the simulated and real data sets are similar to some extent. This file aims to understand their differences. In all figures presented in this supplemental file, red lines correspond to the time profiles of individual subjects, and the black line is the median time profile of that specific group of subjects.

Figure A.1 and A.2 show the time profiles of the subjects with large coefficients in the first component (comp1) in the 3-component CP model for the real data set. These two figures indicate that comp1 (patterns for the real data) mainly captures the subjects with the highest peak appearing around 0.5h in metabolites Pyr, Lac, and Ala. Comparing Figure A.1 and A.2 with Figure A.5, we can see that the time points for the highest peak of Pyr in the real and simulated data are different, leading to the shift of the time point for the highest peak in comp1 for the real vs. simulated data observed in Figure 4 in the main text.

Figure A.3 and A.4 present the time profiles of the subjects with large coefficients in the second component (comp2) in the 3-component CP model for the real data set. From these two figures, we can see that the concentration of Pyr in the selected subjects (subjects with large absolute coefficients in comp2 in the 3-component model for the real data), especially subjects with negative large absolute coefficients (i.e., Figure A.4), possess similar dynamic profiles as Ins and Glc, i.e., dynamic patterns with the highest peak appearing at around 1.5h. However, in the simulated data, Pyr has rather different dynamic profiles compared with Ins and Glc (see Figure A.5). These observations explain why Pyr has different coefficients in comp2 for the real and simulated data sets.

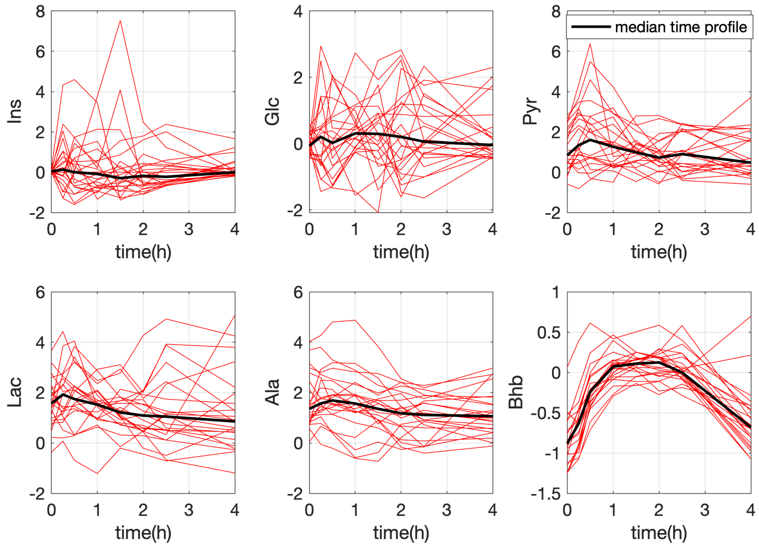


Figure A.1: **Time profiles of subjects with large positive coefficients (no less than 0.08) in comp1 from the 3-component model for the preprocessed real data.**

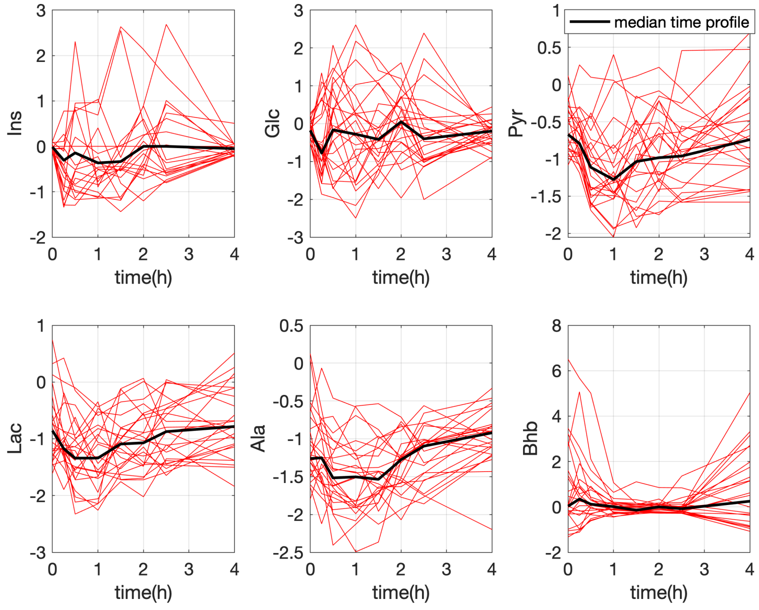


Figure A.2: **Time profiles of subjects with negative large absolute coefficients (no greater than -0.08) in comp1 from the 3-component model for the preprocessed real data.**

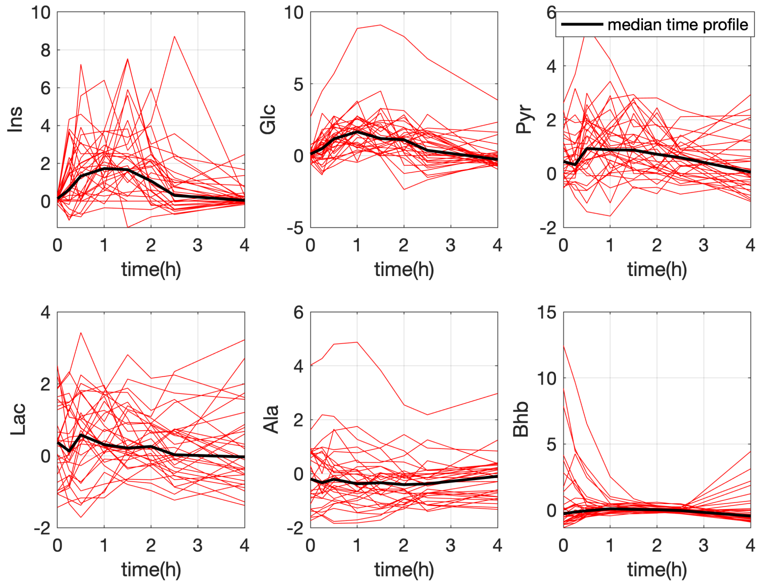


Figure A.3: **Time profiles of subjects with large positive coefficients (no less than 0.08) in comp2 from the 3-component model for the preprocessed real data.**

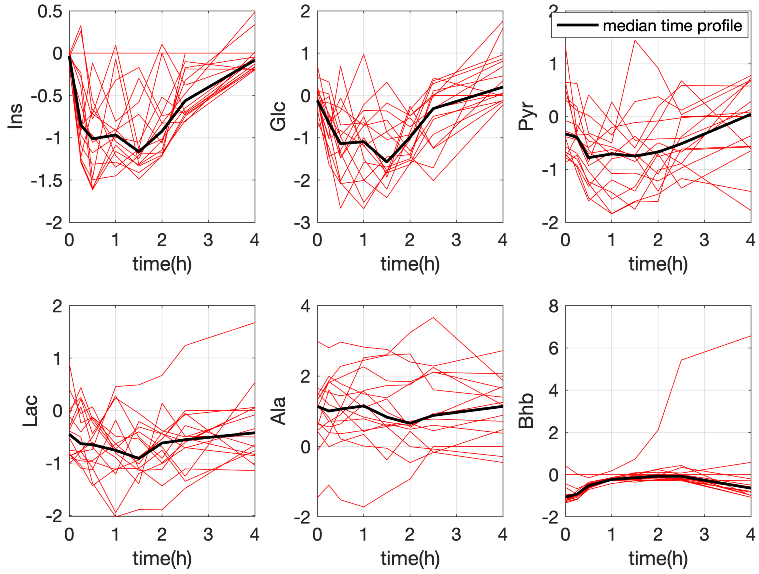


Figure A.4: **Time profiles of subjects with negative large absolute coefficients (no greater than -0.08) in comp2 from the 3-component model for the preprocessed real data.**

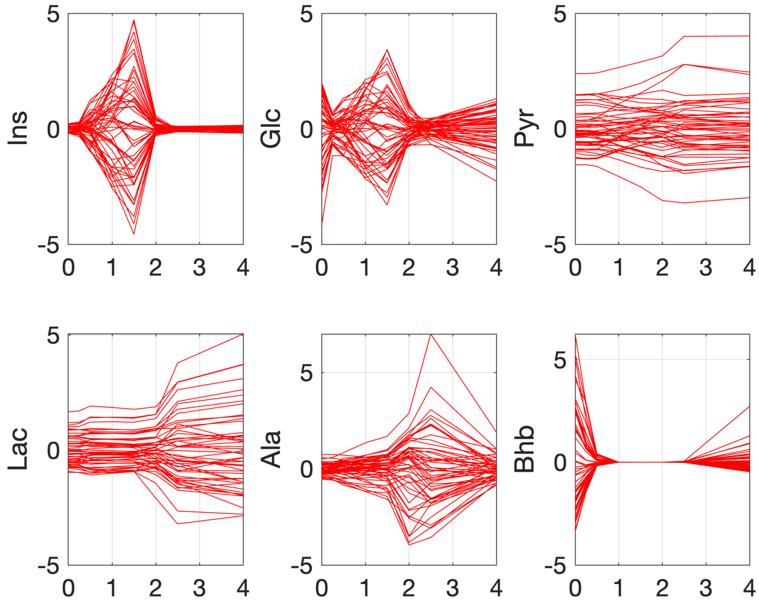


Figure A.5: **Time profiles of each subject for the preprocessed simulated data.**
