## Additional file 5 for "Analyzing postprandial metabolomics data using multiway models: A simulation study"

### Experiments on data sets with random noise

For the simulated three-way data $\mathscr{X} \in \mathbb{R}^{I\times J\times K}$, we add noise to mimic the measurement error by incorporating entries drawn from a standard normal distribution as follows:

$$\mathscr{X}_{\text{noisy}}(:,j,k) = \mathscr{X}(:,j,k) + \eta g \frac{\|\mathscr{X}(:,j,k)\|}{\|g\|},$$

where each element $\mathscr{X}(i,j,k)$ refers to the simulated concentration of the $j$th metabolite for the $i$th subject at the $k$th time point, $\|\cdot\|$ is the Euclidean norm for a vector, $g$ is the vector with length $I$ generated from the standard normal distribution, and $\eta$ is the relative standard deviation (RSD) of a replicate measurement. We consider $\eta$ from 4% to 15%, which aligns with the range mentioned in [1]. We observed negative concentrations of Bhb for some subjects at certain time points with specific values of $\eta$, and we set those negative values to be 0 so that the generated simulated concentrations are meaningful. Our experiments show that the CP models exhibit considerable robustness, particularly when the RSD remains low. For example, when the RSD is 10% or 6% for the data set with *insulin-resistant* vs. control groups or the data set with *beta-cell dysfunction* vs. control groups, respectively, we observe FMS values (noisy vs. noiseless patterns for both the *full-dynamic* and the *T0-corrected* data) greater than 0.92. With an increasing RSD, the added noise may affect the number of components of the CP model and the extracted patterns. In this file, we consider $\eta = 15\%$ and compare the factors extracted by the CP models for the noisy vs. noiseless data.

#### The noisy data set with *insulin-resistant* vs. control group and low within-group variation ($\alpha = 0.2$)

For the *T0-corrected* analysis, we opt for the 3-component CP model illustrated in Figure A.1, rather than the 4-component model employed in the noiseless scenario. This is because the fourth (extra) component in the 4-component CP model captures the random noise instead of a meaningful pattern in the noisy scenario, as demonstrated by the comparison of the 4-component CP models extracted from the noisy versus noiseless *T0-corrected* data in Figure A.2. Furthermore, our findings reveal that the inclusion of noise hinders the replicability of the 4-component CP model. Comparing the analyses for the *T0-corrected* noisy versus noiseless data, as presented in Figure A.3, we observe that the component primarily capturing Pyr and yielding the best subject group separation is not preserved in the noisy scenario; however, this component is merged into the first component giving the best subject group separation.

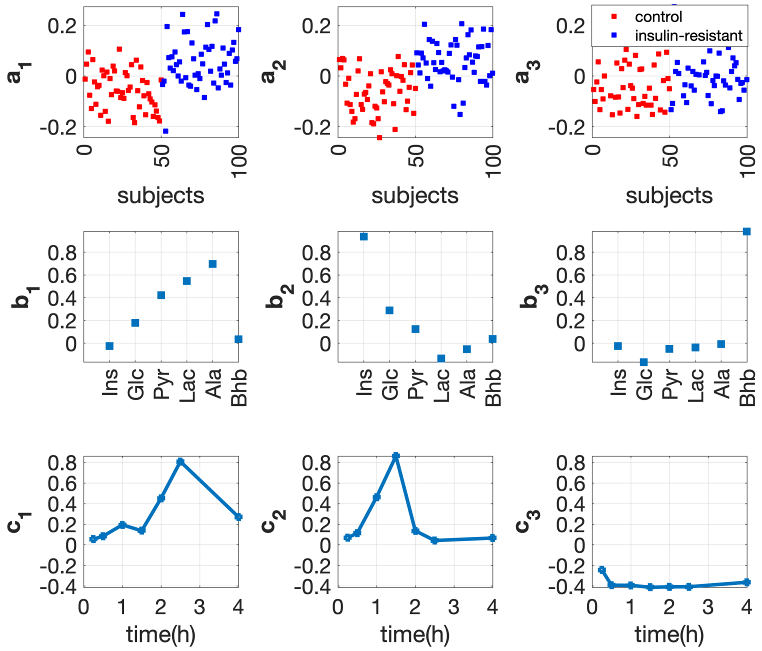


Figure A.1: **Factors of the 3-component CP model extracted from the *T0-corrected* noisy data with *insulin-resistant* and control groups. The individual variation is set by using the random perturbation level $\alpha = 0.2$. The vectors $\mathbf{a}_r$, $\mathbf{b}_r$ and $\mathbf{c}_r$, $r = 1, \cdots, 3$, are the components in the *subjects*, *metabolites* and *time* modes extracted by the CP model.**

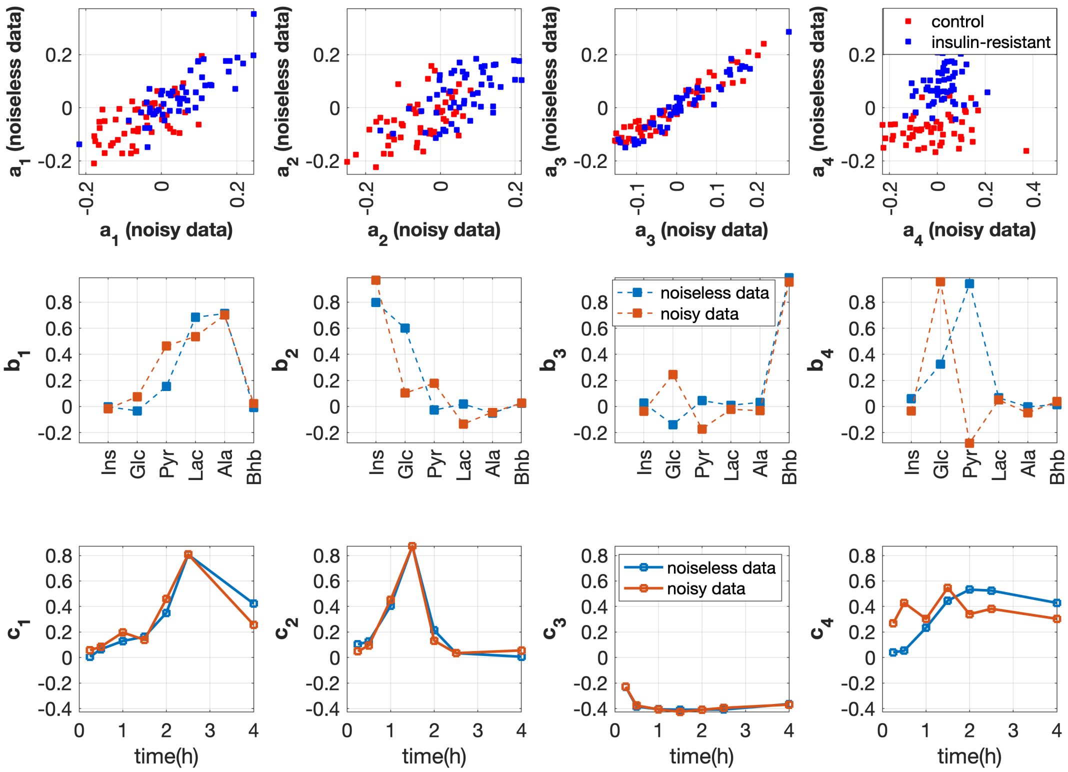


Figure A.2: **Comparison of the 4-component CP models extracted from the *T0-corrected* noisy vs. noiseless data. The data contains 50 *insulin-resistant* and 50 control subjects. The individual variation is set by using the random perturbation level $\alpha = 0.2$. The vectors $\mathbf{a}_r$, $\mathbf{b}_r$ and $\mathbf{c}_r$, $r = 1, \cdots, 4$, are the components in the *subjects*, *metabolites* and *time* modes extracted by the CP model.**

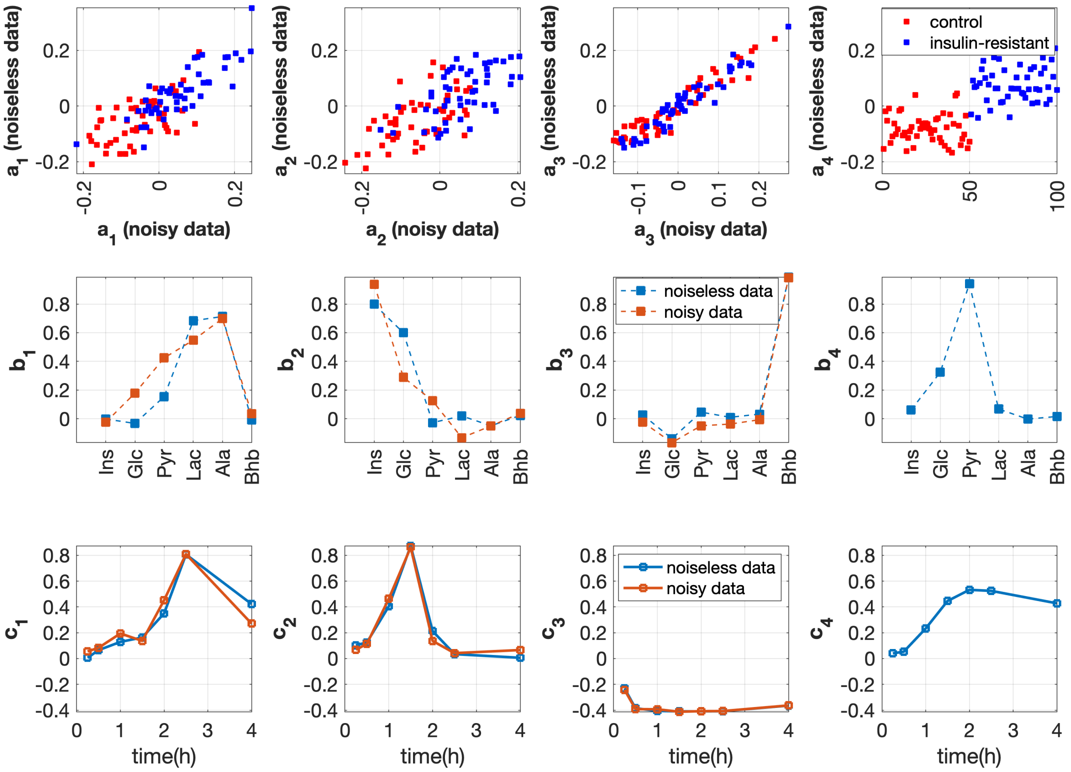


Figure A.3: **Comparison of the factors of the 3-component CP model extracted from the *T0-corrected* noisy data vs. the 4-component model extracted from the noiseless data. The data contains 50 *insulin-resistant* and 50 control subjects. The individual variation is set by using the random perturbation level $\alpha = 0.2$. The vectors $\mathbf{a}_r$, $\mathbf{b}_r$ and $\mathbf{c}_r$, $r = 1, \cdots, 3$, are the components in the *subjects*, *metabolites* and *time* modes extracted by the CP model.**

For the *full-dynamic* analysis, we utilize the 4-component CP model and the factors are presented in Figure A.4. Comparing the analyses for the noisy vs. noiseless data in Figure A.5, we observe that in the presence of noise, the component primarily capturing fasting Pyr is no longer discernible.

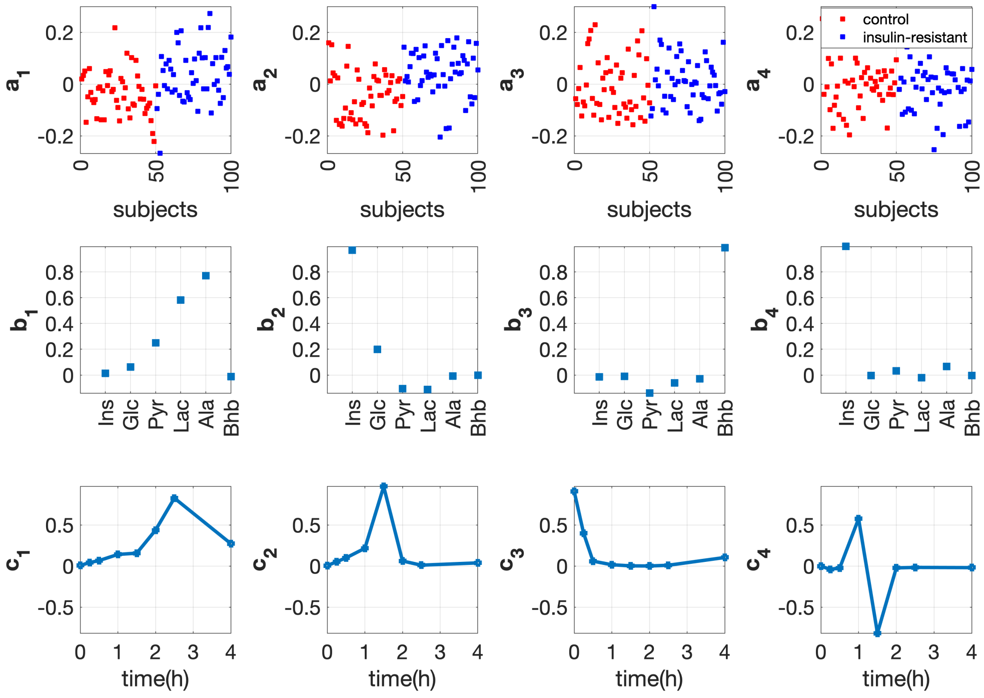


Figure A.4: **Factors of the 4-component CP model extracted from the *full-dynamic* noisy data with *insulin-resistant* and control groups. The individual variation is set by using the random perturbation level $\alpha = 0.2$. The vectors $\mathbf{a}_r$, $\mathbf{b}_r$ and $\mathbf{c}_r$, $r = 1, \cdots, 4$, are the components in the *subjects*, *metabolites* and *time* modes extracted by the CP model.**

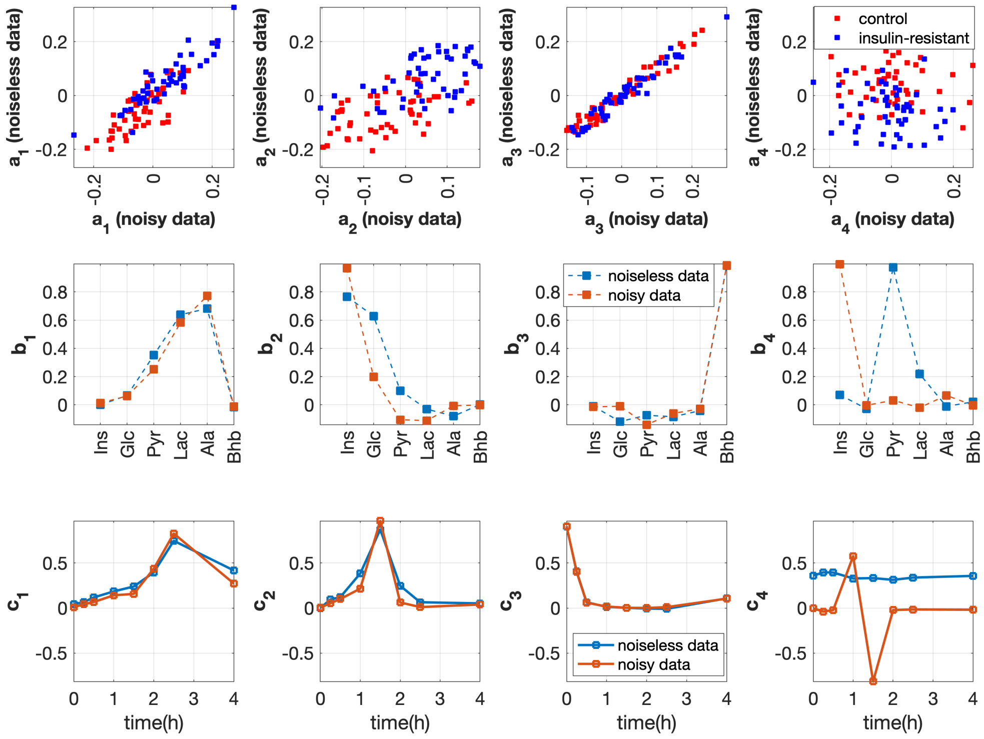


Figure A.5: **Comparison of the factors of the 4-component CP models extracted from the *full-dynamic* noisy vs. noiseless data. The data contains 50 *insulian-resistant* and 50 control subjects. The individual variation is set by using the random perturbation level $\alpha = 0.2$. The vectors $\mathbf{a}_r$, $\mathbf{b}_r$ and $\mathbf{c}_r$, $r = 1, \cdots, 4$, are the components in the *subjects*, *metabolites* and *time* modes extracted by the CP model.**

Comparing Figure A.1 with Figure A.4, we observe that the *T0-corrected* analysis captures the subject group differences better than the *full-dynamic* analysis. This observation aligns with the findings from the analyses of the noiseless data.

#### The noisy data set with *beta-cell dysfunction* vs. control group with low within-group variation ($\alpha = 0.2$)

For the *T0-corrected* noisy data, we employ the 3-component CP model as illustrated in Figure A.6. This choice is made because the additional (fourth) component in the model is influenced by the random noise, resulting in an uninterpretable pattern where Glc and Pyr exhibit opposite score values (refer to Figure A.7 for the comparison of the 4-component CP models from the noisy vs. noiseless data). In addition, the 4-component CP model can not be replicated. Note that we need to remove an outlier in the 4-component CP model when analyzing the *T0-corrected* noisy data; see [2] for details about outlier detections of CP models. From Figure A.8, the comparison of the *T0-corrected* analyses for the noisy vs. noiseless data, we observe that the component primarily capturing individual variations in Glc and Pyr (the fourth component in the noiseless case) disappear when noise is added to the data. However, the component responsible for distinguishing between subject groups shows to be more robust to noise.

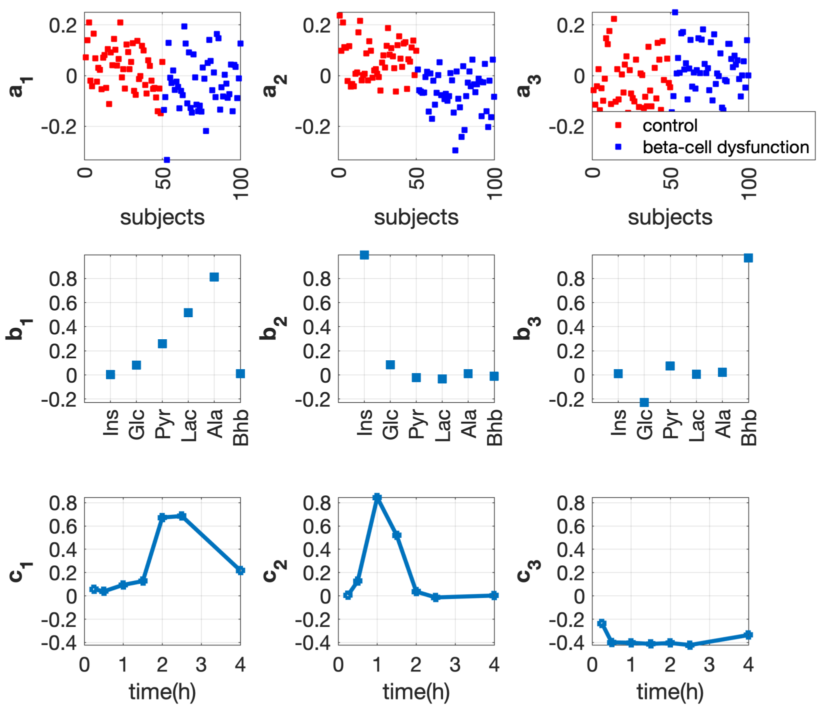


Figure A.6: **Factors of the 3-component CP model extracted from the *T0-corrected* noisy data with *beta-cell dysfunction* and control groups. The individual variation is set by using the random perturbation level as $\alpha = 0.2$. The vectors $\mathbf{a}_r$, $\mathbf{b}_r$ and $\mathbf{c}_r$, $r = 1, \cdots, 3$, are the components in the *subjects*, *metabolites* and *time* modes extracted by the CP model.**

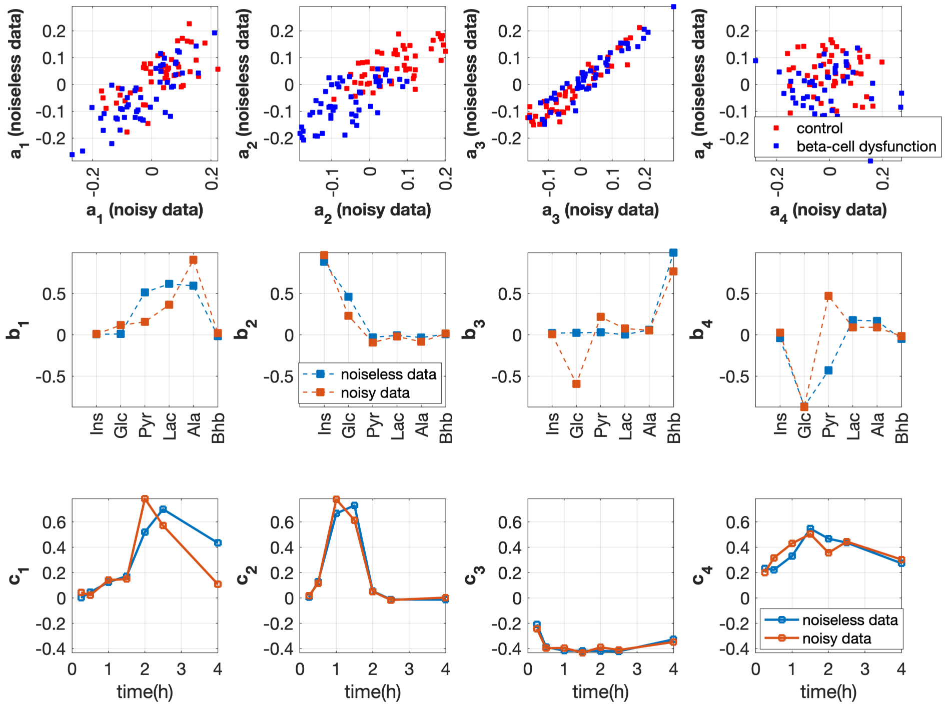


Figure A.7: **Comparison of the factors of the 4-component CP models extracted from the *T0-corrected* noisy vs. noiseless data. The data contains 50 *beta-cell dysfunction* and 50 control subjects. The individual variation is set by using the random perturbation level $\alpha = 0.2$. In this modle, we remove one subject since it is detected as an outlier. The vectors $\mathbf{a}_r$, $\mathbf{b}_r$ and $\mathbf{c}_r$, $r = 1, \cdots, 4$, are the components in the *subjects*, *metabolites* and *time* modes extracted by the CP model.**

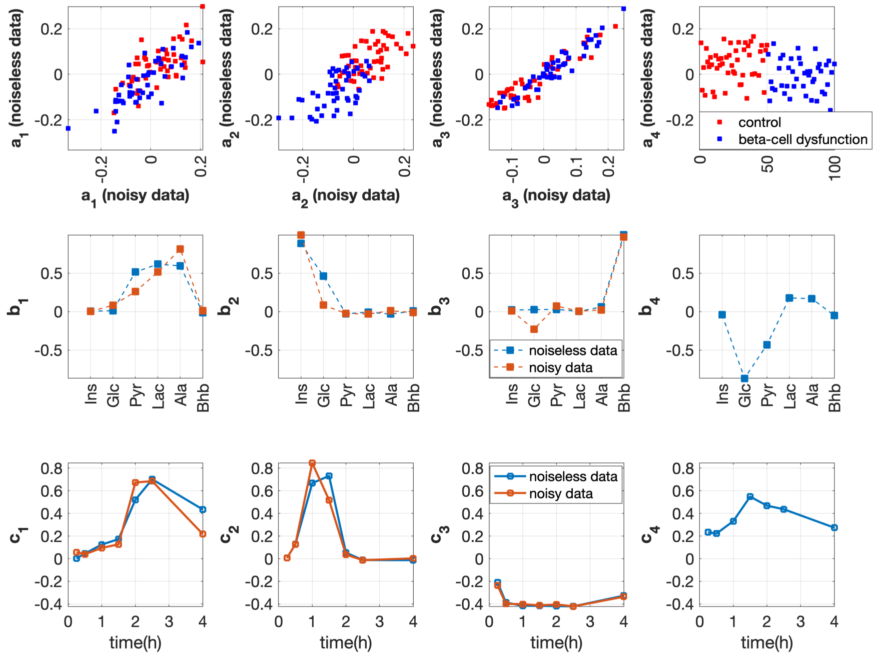


Figure A.8: **Comparison of the factors of the 3-component CP model extracted from the *T0-corrected* noisy data vs. the 4-component model extracted from the noiseless data. The data contains 50 *beta-cell dysfunction* and 50 control subjects. The individual variation is set by using the random perturbation level $\alpha = 0.2$. The vectors $\mathbf{a}_r$, $\mathbf{b}_r$ and $\mathbf{c}_r$, $r = 1, \cdots, 4$, are the components in the *subjects*, *metabolites* and *time* modes extracted by the CP model.**

For the *full-dynamic* data, we employ the 4-component CP model, and the corresponding factors are presented in Figure A.9. Comparing the analyses of the noisy with the noiseless data in Figure A.10, we note that despite the presence of noise, the CP model remains capable of capturing the differences between subject groups and the corresponding significant metabolites (the second and fourth components). However, due to the noise, the scores associated with these significant metabolites vary, resulting in a slight shift in the peak values within the dynamic profiles.

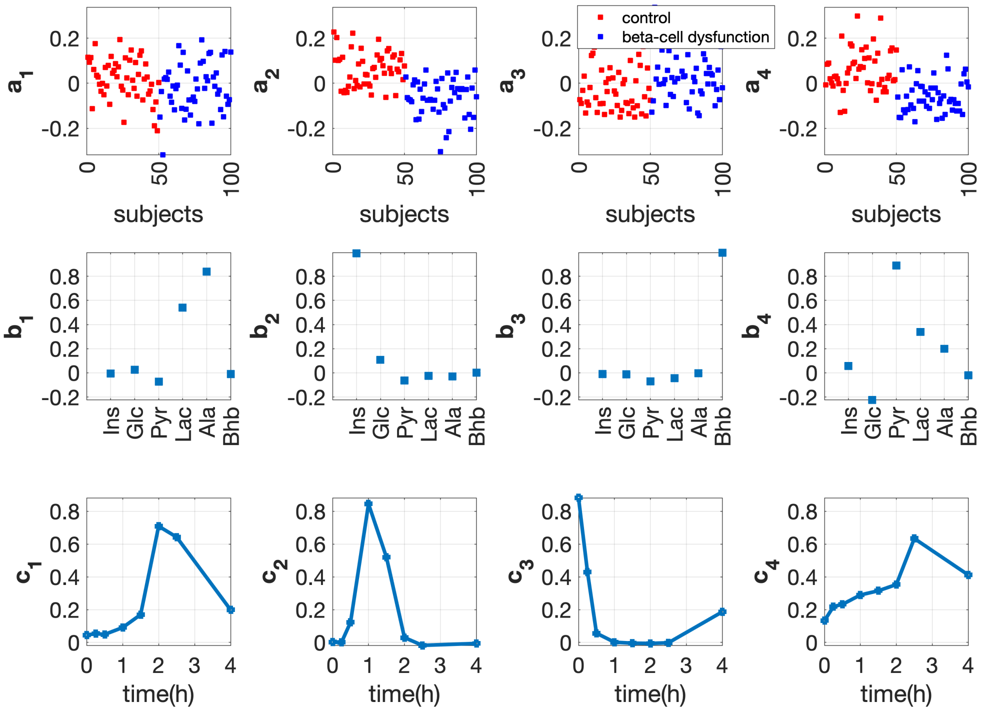


Figure A.9: **Factors of the 4-component CP model extracted from the *full-dynamic* noisy data with *beta-cell dysfunction* and control groups. The individual variation is set by using the random perturbation level $\alpha = 0.2$. The vectors $\mathbf{a}_r$, $\mathbf{b}_r$ and $\mathbf{c}_r$, $r = 1, \cdots, 4$, are the components in the *subjects*, *metabolites* and *time* modes extracted by the CP model.**

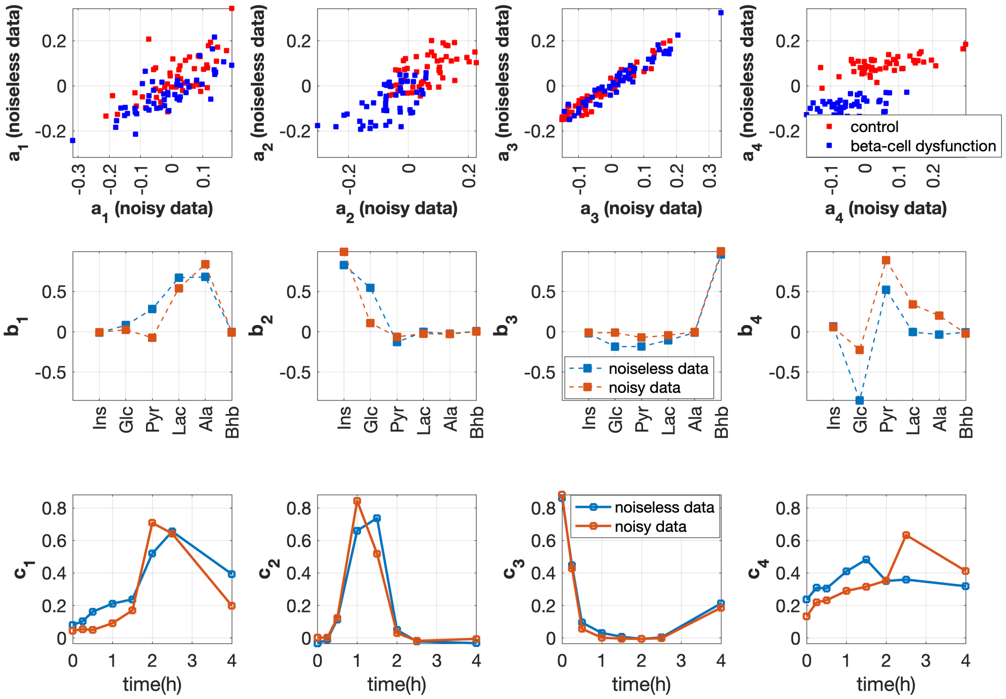


Figure A.10: **Comparison of the factors of the 4-component CP models extracted from the *full-dynamic* noisy data vs. data without noise. The data contains 50 *beta-cell dysfunction* and 50 control subjects. The individual variation is set by using the random perturbation level $\alpha = 0.2$. The vectors $\mathbf{a}_r$, $\mathbf{b}_r$ and $\mathbf{c}_r$, $r = 1, \cdots, 4$, are the components in the *subjects*, *metabolites* and *time* modes extracted by the CP model.**

### References

[1] Age K Smilde, Mariët J van der Werf, Jean-Pierre Schaller, and Cor Kistemaker. Characterizing the precision of mass-spectrometry-based metabolic profiling platforms. *Analyst*, 134(11):2281–2285, 2009.

[2] Rasmus Bro. PARAFAC. Tutorial and applications. *Chemometrics and Intelligent Laboratory Systems*, 38(2):149–171, 1997.
